# Interneuronal mechanisms of hippocampal theta oscillation in full-scale model of the CA1 circuit

**DOI:** 10.1101/087403

**Authors:** Marianne J. Bezaire, Ivan Raikov, Kelly Burk, Dhrumil Vyas, Ivan Soltesz

## Abstract

The hippocampal theta rhythm plays important roles in information processing; however, the mechanisms of its generation are not well understood. We developed a data-driven, supercomputer-based, full-scale (1:1) model of the CA1 area and studied its interneurons during theta oscillations. Theta rhythm with phase-locked gamma oscillations and phase-preferential discharges of distinct in terneuronal types spontaneously emerged from the isolated CA1 circuit without rhythmic inputs. Perturbation experiments identified parvalbumin-expressing interneurons and neurogliaform cells, as well as interneuronal diversity itself, as important factors in theta generation. These simulations reveal new insights into the spatiotemporal organization of the CA1 circuit during theta oscillations.

## Introduction

The hippocampal CA1 area supports diverse cognitive tasks including learning, memory, and spatial processing (Squire, 1992; Remondes and Schuman, 2004; Manns et al., 2007; Moser et al., 2008). These cognitive tasks are thought to require coordination of neuronal activity provided by physiological network oscillations, including the theta rhythm (Buzsáki, 2002; Buzsáki and Moser, 2013). In rodents, hippocampal theta is a 5-10 Hz oscillation in the local field potential (LFP) and neuronal firing probabilities (Soltesz and Deschenes, 1993; Lee et al., 1994; Ylinen et al., 1995; Klausberger and Somogyi, 2008; Varga et al., 2012, 2014), occurring during locomotion and in REM sleep (Buzsáki, 2002).

Though several major afferents provide theta-frequency rhythmic input to the CA1 *in vivo* (Soltesz and Deschenes, 1993; Buzsáki, 2002; Fuhrmann et al., 2015), recent reports indicate the presence of spontaneous theta-frequency LFP oscillations even in the isolated whole CA1 preparation *in vitro* (Goutagny et al., 2009; Amilhon et al., 2015). Therefore, the latter studies suggest an intrinsic ability of the CA1 circuit to generate some form of theta waves even without rhythmic external inputs. However, the intra-CA1 mechanisms that may contribute to the generation of the theta rhythm are not well understood (Colgin, 2013, 2016).

Here we investigated the ability of the CA1 to generate intrinsic theta oscillations using a uniquely biological data-driven, full-scale computer model of the isolated CA1 network. Recent advances in supercomputing power and high-quality synaptic connectivity data present the intriguing opportunity to develop full-scale models where every biological synapse and neuron is explicitly represented. In principle, such full-scale models of mammalian circuits comprising hundreds of thousands of neurons of distinct types advantageously avoid the connectivity scaling tradeoff that besets reduced-scale models: smaller models of large networks with realistic single cell electrophysiological properties (e.g., input resistance and resting membrane potential) remain silent unless synaptic strengths or numbers are arbitrarily increased beyond the biologically relevant levels to compensate for fewer inputs to their model cells (e.g., Dyhrfjeld-Johnsen et al. (2007); Sterratt et al. (2011)). Biological relevance may also increase as other network components are modeled in greater detail. However, full-scale models require considerable computational resources. Further, such detailed models have a large parameter space which risks being sub-optimally constrained by neurobiological properties that are only partially quantified (Sejnowski et al., 1988). Because the CA1 area is one of the most extensively studied brain regions, there are abundant anatomical and electrophysiological data about its organization, making it a logical choice for the development of a full-scale model. The CA1 area is also worth modeling at full-scale because of the diverse cognitive tasks it supports. These tasks likely require the simultaneous processing of thousands of incoming and outgoing signals, and full-scale network models, at least in principle, have the potential to match this *in vivo* processing capacity.

In this paper, we describe the development of a full-scale CA1 computational network model of unprecedented biological detail and its application to gain insights into the roles and temporal organization of CA1 interneurons during theta rhythm. The simulated full-scale CA1 circuit was able to spontaneously generate theta waves as well as phase-locked gamma oscillations. Furthermore, distinct interneuron types discharged at particular phases of theta, demonstrating that phase-preferential firing (Klausberger et al., 2003, 2004, 2005; Ferraguti et al., 2005; Jinno et al., 2007; Fuentealba et al., 2008; Klausberger and Somogyi, 2008; Varga et al., 2012; Lapray et al., 2012; Katona et al., 2014; Varga et al., 2014) originates in part within the CA1 network. Perturbation experiments revealed that parvalbumin-expressing (PV+) interneurons, neurogliaform cells, connections beween CA1 pyramidal cells, and interneuronal diversity were important for theta generation. These results provide new mechanistic insights into the emergence of the theta rhythm from within the CA1 circuitry and the role of interneurons in theta oscillations.

## Results

### Development of data-driven, full-scale model of the isolated CA1

Details of the full-scale model are described in the Methods, and the most important features are illustrated in Figures 1 and 2 and summarized here. Briefly, CA1 model cells were evenly distributed within their respective layers in a 3-dimensional prism with realistic dimensions for the rodent hippocampal CA1 region (Figure 1A and 1B). The model network contained 338,740 cells (similar to the biological CA1 in rats, including 311,500 pyramidal cells and 27,240 interneurons) (Figure 1D-1E and Figure 1 - figure supplement 1). In addition, the network also incorporated 454,700 artificial stimulating cells (spiking units with random, Poisson-distributed inter-spike intervals) to simulate afferents to CA1; the cell type-specific distribution, dendritic position, amplitude and kinetics of the excitatory input synapses were all experimentally constrained by afferent CA3 and entorhinal cortical data. Cell type-specific connectivity data, including cell numbers (Figure 1D) and convergence and divergence values (Figure 1E; Figure 1 - figure supplement 1 and Table 1) were taken without alteration from our previously published, in-depth, quantitative assessment of the CA1 circuit (Bezaire and Soltesz, 2013). Anatomical constraints of the connectivity were implemented in the model by accounting for the distribution of the axonal boutons as a function of longitudinal and transverse distance from the presynaptic cell soma (Figure 1 - figure supplement 2). The afferent divergence and convergence onto the cells were also anatomically patterned, maintaining the topographical arrangement seen experimentally (Hongo et al., 2015), for a total of 5.19 billion synaptic connections in the model network. In addition, the remaining parameters that could not be constrained by experimental data were documented, with the assumptions used to arrive at them explicitly listed in Table 2 of Bezaire and Soltesz (2013) and additional parameter calculations described in the Supplementary Material Section 3 of the present paper. To highlight the many constraints applied in the current work and address the unconstrained model parameters, we characterized all model components (constrained and unconstrained) in experimental terms, comparing with experimental data where possible (Figure 2; Supplementary Material). For a four second simulation, the full-scale model required 3-4 terabytes (TB) of RAM and four hours of execution time on a supercomputer using ~3000 processors (or up to 12 hours for simulations calculating a high-accuracy local field potential (LFP) analog). Additional details and data about model performance are available in Table 2 and Bezaire et al. (2016).

**Figure 1:**
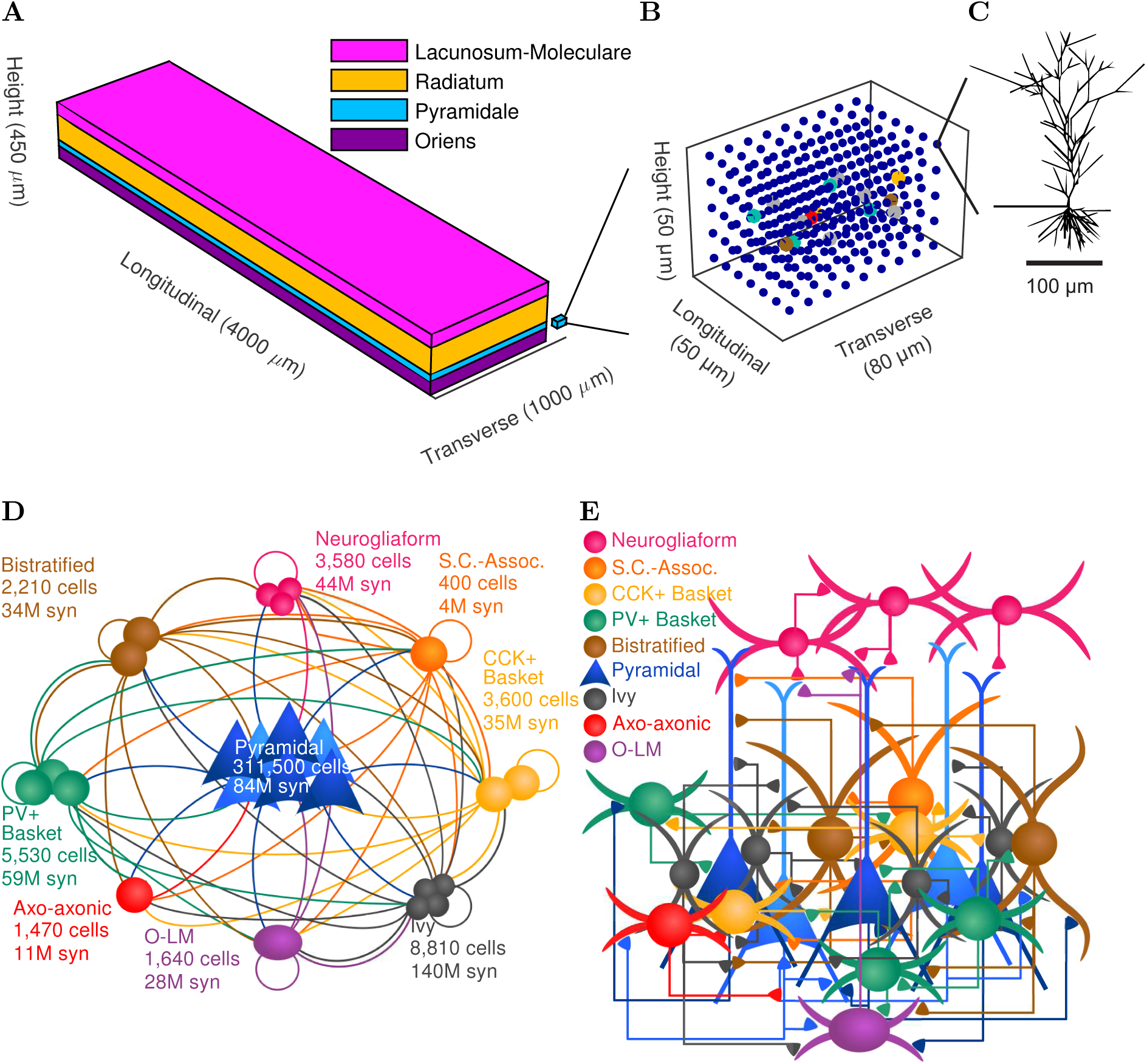
CA1 network connectivity. (A) The model network is arranged in a layered prism with the lengths of each dimension similar to the actual dimensions of the CA1 region and its layers. (B) The model cell somata within a small chunk of stratum pyramidale (as depicted in A) are plotted to show the regular distribution of model cells throughout the layer in which they are found. (C) Each pyramidal cell in the network has detailed morphology with realistic incoming synapse placement along the dendrites and soma. (D-E) Diagrams illustrate connectivity between types of cells. (D) The network includes one principal cell type (pyramidal cells) and eight interneuron types. Cell types that may connect are linked by a line colored according to the presynaptic cell type. Most cell types can connect to most other cell types. Total number of cells of each type are displayed, as are the number of local output synapses (boutons) from all cells of each type. (E) The number, position, and cell types of each connection are biologically constrained, as are the numbers and positions of the cells. See Figure 1 - figure supplement 1 for details about the convergence onto each cell type. Also see Table 1 and Figure 1 - figure supplement 2 for information about the cell-type combinations of the 5 billion connections and the axonal distributions followed by each cell type, as well as detailed connectivity results at http://doi.org/10.6080/K05H7D60.

**Figure 2:**
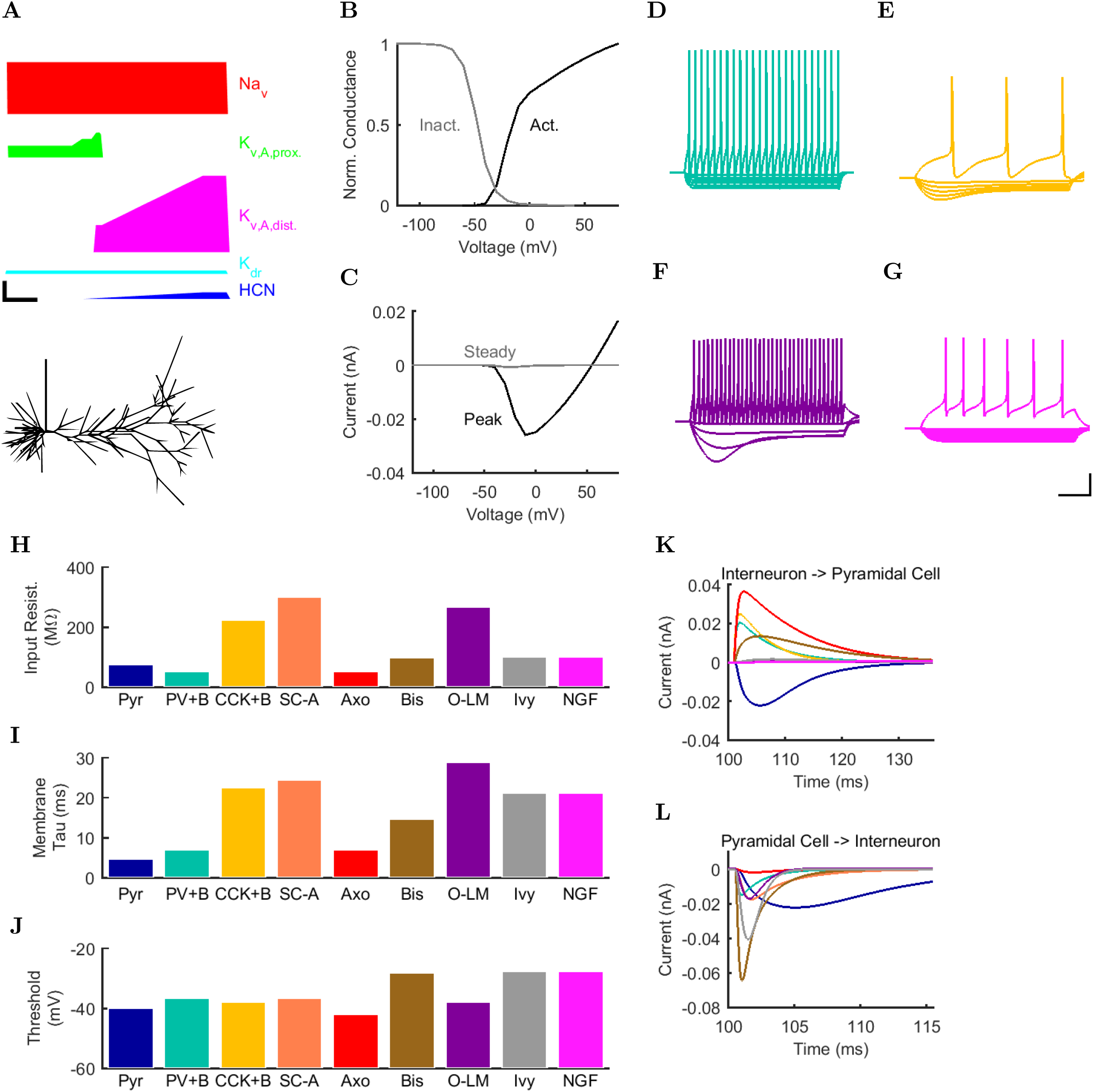
Electrophysiology of the model network components. (A) Ion channel densities vary as a function of location (top) in the morphologically detailed pyramidal cell model (bottom; adapted from Poolos et al. (2002)). Scale bar: 100 *µ*m and 0.01 *µ*F/cm^2^. (B - C) The sodium channel found in the pyramidal cell soma is characterized in terms of (B) the activation/inactivation curves and (C) the current-voltage relation at peak (transient) current and steady state. (D - (G)) Current sweeps are shown for 4 model cell types: (D) PV+ basket cell, (E) CCK+ basket cell, (F) O-LM cell, and (G) neurogliaform cell. Scale bar: 100 ms and 20 mV. (H-J) Electrophysiological properties for each cell type, including (H) input resistance, (I) membrane time constant, and (J) action potential threshold. (K - L) Pyramidal cell synaptic connections are characterized as post-synaptic currents with the postsynaptic cell voltage clamped at -50 mV; (K) synapses made onto the pyramidal cell from all other cell types and (L) synapses made by the pyramidal cell onto all network cell types. Cells represented by same colors as in Figure 1. Source Data available at Figure 2 - Source Data.zip. Additional details available in the Methods, Table 3, and the Supplementary Material.

**Table 1:** Number of synapses between each cell type. Connections between cells generally comprise 1 - 10 synapses each. Presynaptic cells are listed down the first column (corresponding to each row) and postsynaptic cells are listed along the first row (corresponding to each column).

| **Pre/Post** | **Axo** | **Bis** | **CCK+B** | **Ivy** | **NGF** | **O-LM** | **Pyr** | **PV+B** | **SC-A** |
| --- | --- | --- | --- | --- | --- | --- | --- | --- | --- |
| Axo | 0.00e+00 | 0.00e+00 | 0.00e+00 | 0.00e+00 | 0.00e+00 | 0.00e+00 | 1.12e+07 | 0.00e+00 | 0.00e+00 |
| Bis | 2.35e+05 | 3.54e+05 | 5.76e+05 | 2.64e+05 | 0.00e+00 | 6.40e+05 | 3.12e+07 | 8.85e+05 | 6.80e+04 |
| CCK+B | 1.41e+05 | 2.12e+05 | 9.79e+05 | 5.64e+05 | 0.00e+00 | 2.62e+05 | 3.24e+07 | 5.31e+05 | 8.32e+04 |
| Ivy | 3.53e+05 | 5.30e+05 | 3.42e+06 | 2.11e+06 | 1.00e+06 | 2.23e+06 | 1.28e+08 | 1.33e+06 | 4.08e+05 |
| NGF | 0.00e+00 | 0.00e+00 | 0.00e+00 | 0.00e+00 | 6.09e+05 | 0.00e+00 | 4.36e+07 | 0.00e+00 | 0.00e+00 |
| O-LM | 1.18e+05 | 1.77e+05 | 1.44e+06 | 0.00e+00 | 4.65e+05 | 9.84e+04 | 2.49e+07 | 4.42e+05 | 1.60e+05 |
| Pyr | 7.19e+05 | 2.43e+06 | 0.00e+00 | 2.38e+05 | 0.00e+00 | 1.17e+07 | 6.14e+07 | 7.03e+06 | 1.26e+05 |
| PV+B | 5.73e+04 | 8.62e+04 | 1.37e+05 | 7.05e+04 | 0.00e+00 | 0.00e+00 | 5.83e+07 | 2.16e+05 | 9.60e+03 |
| SC-A | 8.82e+03 | 1.33e+04 | 1.30e+05 | 1.06e+05 | 0.00e+00 | 1.97e+04 | 3.74e+06 | 3.32e+04 | 1.44e+04 |
| CA3 | 1.23e+07 | 2.56e+07 | 1.44e+07 | 3.39e+07 | 0.00e+00 | 0.00e+00 | 3.73e+09 | 6.69e+07 | 1.55e+06 |
| ECIII | 1.43e+06 | 1.91e+06 | 4.02e+06 | 0.00e+00 | 3.75e+06 | 0.00e+00 | 8.09e+08 | 0.00e+00 | 4.58e+05 |

**Table 2:** Simulation time, exchange time, and load balance for simulations executed on various supercomputers and numbers of processors.

| **Supercomputer** | **# Processors** | **Sim Time (s)** | **Exchange Time (s)** | **Load Balance** |
| --- | --- | --- | --- | --- |
| Comet | 1680 | 2610.28 | 1.05 | 0.999 |
| Comet | 1704 | 2566.76 | 0.65 | 0.999 |
| Comet | 1728 | 2601.22 | 0.86 | 0.999 |
| Comet via NSG | 1728 | 2060.88 | 0.83 | 0.999 |
| Stampede via NSG | 2048 | 2471.64 | 1.71 | 1.000 |
| Stampede | 2048 | 2578.32 | 0.29 | 1.000 |
| Stampede | 2528 | 2189.56 | 1.78 | 0.999 |
| Stampede | 3008 | 1844.22 | 0.91 | 0.999 |
| Stampede | 3488 | 1641.91 | 0.86 | 0.999 |

An important set of constraints was the electrophysiology and other properties of individual cells and synapses (Figure 2; Tables 3 and 4) that were based on experimental data. Briefly, our pyramidal cell model (Poolos et al., 2002) contained 200 compartments in a realistic morphology and six fully characterized ion channel types with kinetics and densities based on anatomical location within the cell (Figure 2A-2C). We included eight model interneuron types (Klausberger and Somogyi, 2008; Soltesz, 2006; Armstrong and Soltesz, 2012): PV+ basket cells (these fast-spiking cells synapse on the somata and proximal dendrites of CA1 pyramidal cells), cholecystokinin+ (CCK+) basket cells (these regular-spiking cells also innervate the somata and proximal dendrites, but have properties and functions distinct from the PV+ basket cells), bistratified cells (these PV+ and somatostatin+ (SOM+) fast-spiking cells innervate the basal and apical dendritic trees), axo-axonic cells (these PV+ fast-spiking cells exclusively synapse on the axon initial segments of pyramidal cells and are also known as chandelier cells), Schaffer Collateral-Associated (SC-A) cells (these CCK+, regular-spiking cells innervate dendrites in the stratum radiatum), oriens-lacunosum-moleculare (O-LM) cells (these SOM+ cells project to the distal dendrites in the stratum lacunosum-moleculare though their somata are located in the stratum oriens), neurogliaform cells (these cells have relatively small dendrites and a dense axonal cloud, and they innervate distal dendrites in the stratum lacunosum-moleculare), and ivy cells (these cells are similar to neurogliaform cells, but innervate proximal dendrites) (Figure 2D-2J). Some interneurons in the model, as in the biological network, also innervated other interneurons (Table 1). For greater detail of model connectivity, including convergence per single cell, synaptic amplitude, and other factors, see the Supplementary Material. These cell types collectively comprise the majority (~70%) of known CA1 interneurons (Bezaire and Soltesz, 2013). The remaining 30% of the interneurons were not included in the model due to paucity of quantitative data (Bezaire and Soltesz, 2013). We differentiated the interneurons by their electrophysiological profiles, connectivity patterns, synaptic properties, and anatomical abundance (Gulyas et al., 1991; Hajos and Mody, 1997; Maccaferri et al., 2000; Megías et al., 2001; Lee et al., 2010; Krook-Magnuson et al., 2011; Bezaire and Soltesz, 2013; Lee et al., 2014). The synaptic connections were implemented using double exponential mechanisms to better fit experimental data on rise and decay time constants. We used experimental data to constrain the synaptic kinetics, amplitudes, and locations on the postsynaptic cell (Figures 1E, 2K, and 2L). We implemented the model in parallel NEURON (Carnevale and Hines, 2005) and executed the simulations on several supercomputers. All model results, characterizations, and experimental comparisons are publically available.

**Table 3:** Electrophysiological characteristics of each model cell type. For more information about model electrophysiology, see the Supplementary Material.

| **Condition** | **Pyr** | **PV+B** | **CCK+B** | **SC-A** | **Axo** | **Bis** | **O-LM** | **Ivy** | **NGF** |
| --- | --- | --- | --- | --- | --- | --- | --- | --- | --- |
| Resting Membrane Potential (mV) | -63.0 | -65.0 | -70.6 | -70.5 | -65.0 | -67.0 | -71.5 | -60.0 | -60.0 |
| Input Resistance ($M\Omega$) | 62.2 | 52.0 | 211.0 | 272.4 | 52.0 | 98.7 | 343.8 | 100.0 | 100.0 |
| Membrane Tau (ms) | 4.8 | 6.9 | 22.6 | 24.4 | 7.0 | 14.7 | 22.4 | 21.1 | 21.1 |
| Rheobase (pA) | 250.0 | 300.0 | 60.0 | 40.0 | 200.0 | 350.0 | 50.0 | 160.0 | 170.0 |
| Threshold (mV) | 52.0 | -36.6 | -40.6 | -43.1 | -41.6 | -28.1 | 100.2 | -27.6 | -27.7 |
| Delay to 1st Spike (ms) | 12.4 | 74.6 | 166.6 | 127.7 | 43.5 | 28.4 | 8.9 | 173.3 | 119.0 |
| Half-Width (ms) | 80.7 | 0.9 | 1.9 | 1.6 | 0.6 | 0.5 | 112.9 | 0.6 | 0.6 |

**Table 4:** Current injection levels used to characterize interneuron current sweeps in Figure 2D-2G.

| Cell Type | Hyper. (pA) | Step Size (pA) | Depol. (pA) |
| --- | --- | --- | --- |
| PV+ B. | -300 | 50 | +500 |
| CCK+ B. | -100 | 20 | +80 |
| O-LM | -130 | 30 | +80 |
| NGF | -130 | 20 | +190 |

### Emergence of spontaneous theta and gamma oscillations in the full-scale model in the absence of rhythmic external inputs

First, we examined whether the well-constrained, biologically detailed, full-scale CA1 model could oscillate spontaneously within the physiological range. Based on reports of spontaneous theta-frequency LFP oscillations in the isolated CA1 preparation (Goutagny et al., 2009), we expected a sufficiently constrained CA1 model to generate spontaneous theta rhythm when given tonic, arrhythmic excitation. We varied the magnitude of arrhythmic, tonic excitation to the network (by systematically changing the mean spiking frequency of the artificial stimulating cells, see above) and identified excitation levels where the network developed a stable, spontaneous theta rhythm (5-10 Hz; Figures 3 and 4). The pyramidal cell spikes (Figures 3C and 3D) exhibited peak power around the theta frequency of 7.8 Hz (Figure 4 and Table 7). Importantly, every measure of network activity showed theta oscillations, including the somatic intracellular membrane potential from individual cells (Figure 3D), the spike times of individual cells and all cells collectively (Figure 3C), and aggregate measures such as the spike density function (Szűcs, 1998) per cell type and the LFP analog (Figures 3A and 4; see also Figure 4 - figure supplement 1). In all of these measures of network activity, theta was apparent within one theta period of the simulation start. The theta oscillation was stable, maintaining a steady power level throughout the duration of the oscillation (Figure 4A). To our knowledge, this is the first strictly data-driven, full-scale computational network model of the CA1 that exhibits spontaneous theta rhythm without rhythmic synaptic inputs.

**Figure 3:**
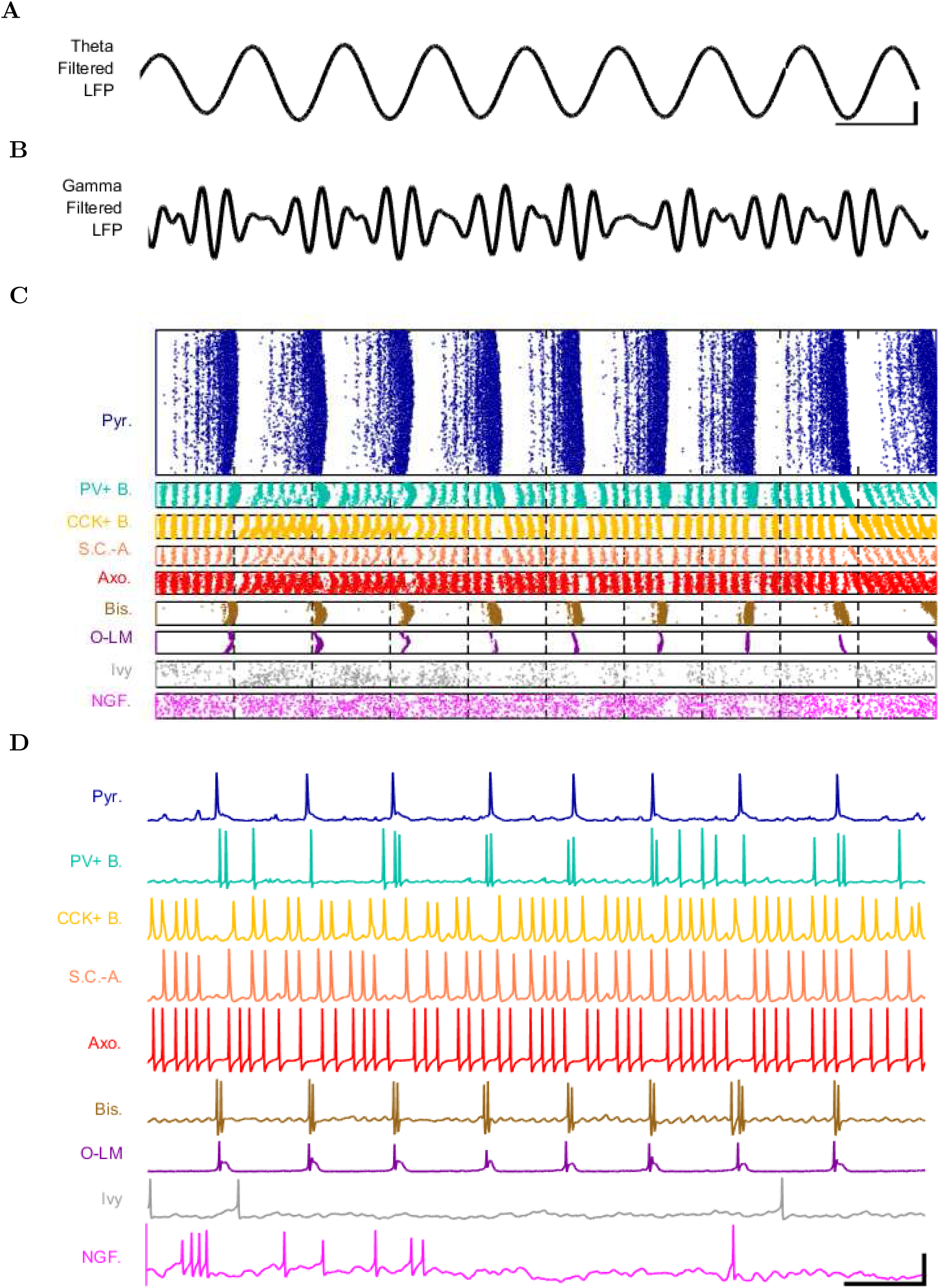
Detailed network activity. (A-D) One second of network activity is shown. (A-B) The LFP analog, filtered at (A) the theta range of 5-10 Hz and (B) the low gamma range of 25-40 Hz, shows consistent theta and gamma signals. Scale bar represents 100 ms and 0.2 mV (theta) or 0.27 mV (gamma) for filtered LFP traces. (C) Raster of all spikes from cells within 100 *µ*m of the reference electrode point. (D) Representative intracellular somatic membrane potential traces from cells near the reference electrode point. Scale bar represents 100 ms and 50 mV for the intracellular traces. Source Data available at Figure 3 - Source Data.zip.

**Figure 4:**
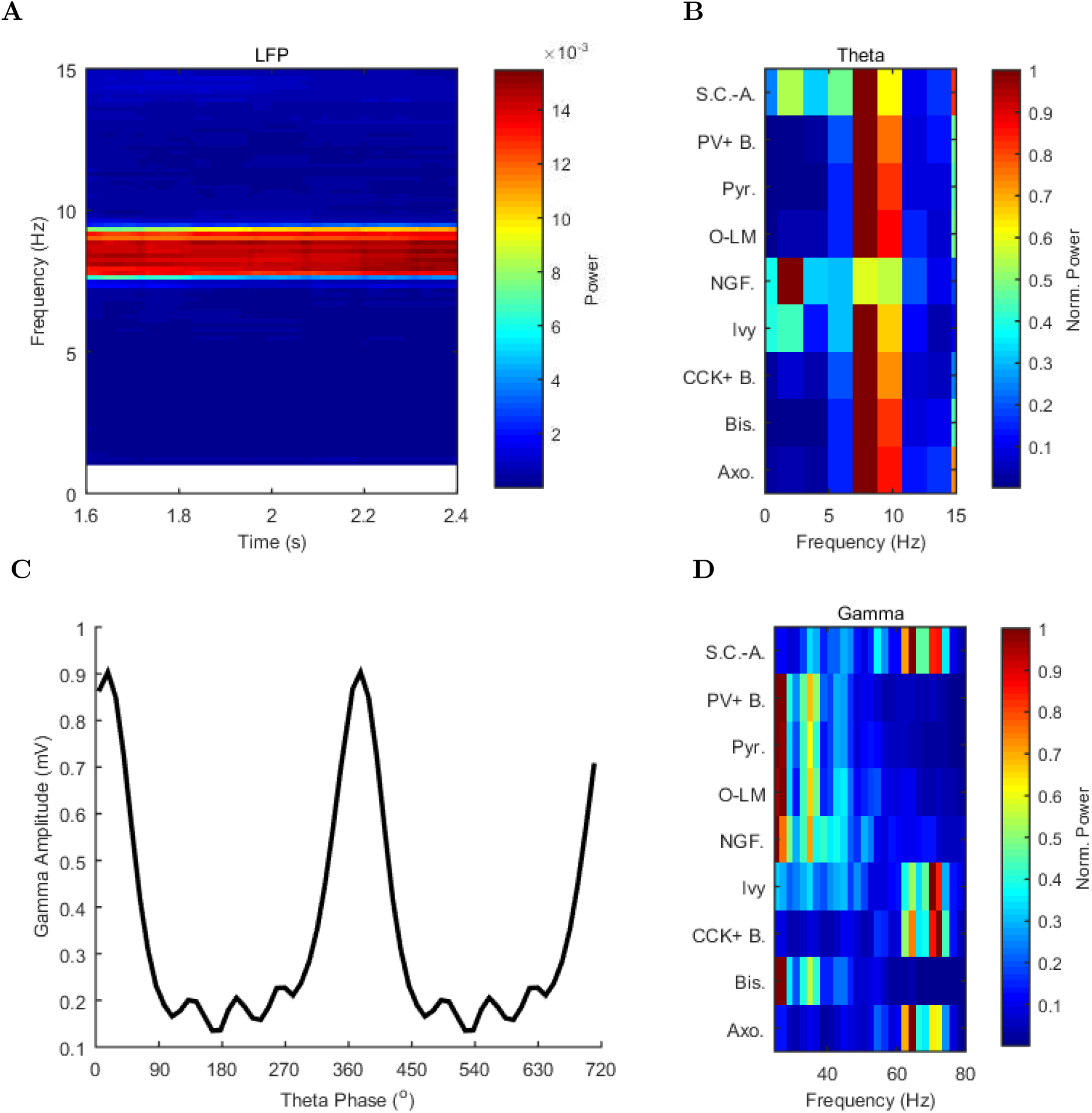
Spectral analysis of model activity. (A) A spectrogram of the local pyramidal-layer LFP analog (including contributions from all pyramidal cells within 100 *µ*m of the reference electrode and 10% of pyramidal cells outside that radius) shows the stability and strength of the theta oscillation over time. The oscillation also featured strong harmonics at multiples of the theta frequency of 7.8 Hz. (B,D) Welch’s periodogram of the spike density function for each cell type, normalized by cell type and by displayed frequency range, shows the dominant network frequencies of (B) theta (7.8 Hz) and (D) gamma (71 Hz). Power is normalized to the peak power displayed in the power spectrum for each cell type. (C) Cross-frequency coupling between theta and gamma components of the LFP analog shows that the gamma oscillation is theta modulated. The gamma envelope is a function of the theta phase with the largest amplitude gamma oscillations occurring at the trough of the theta oscillation. Following convention, the theta trough was designated 0*°*/360*°*; see e.g., Varga et al. (2012). A graphical explanation of the relation between a spike train and its spike density function is shown in Figure 4 - figure supplement 1. Source Data available for this figure at Figure 4 - Source Data.zip.

In addition to theta rhythm, the model network displayed gamma oscillations (25-80 Hz; Figures 3B and 4D), as expected based on *in vivo* data (Soltesz and Deschenes, 1993; Tort et al., 2009; Colgin and Moser, 2010) and *in vitro* slice data showing 65-75 Hz gamma oscillations arising in response to theta rhythmic network stimulation (Butler et al., 2016). The amplitude envelope of the gamma oscillation was phase-locked to the theta rhythm (Figure 3A, 3B and 4C), as it is in the biological CA1, representing cross-frequency coupling (Soltesz and Deschenes, 1993; Bragin et al., 1995; Buzsáki et al., 2003; Jensen and Colgin, 2007; Belluscio et al., 2012). The highest amplitude of the gamma oscillations in the model was observed at the theta trough (0^*°*^/360^*°*^) in the pyramidal layer LFP analog (Figure 4C). Because the current study focused primarily on theta oscillations and experimental data from the isolated CA1 are available only for the theta rhythm (Goutagny et al., 2009; Amilhon et al., 2015), the gamma oscillations were not examined further in the present study.

These results demonstrate that, in spite of gaps in our knowledge, our model was sufficiently well-constrained by experimental data that it generated theta and gamma oscillations on its own, without extrinsic rhythmic inputs or deliberate tuning of intrinsic parameters.

Although in this paper we generally refrained from deliberately compensating for missing parameters, it is of course possible to do so. For example, as mentioned above, no sufficiently detailed information was available for certain interneuron types. Therefore, these lesser-known interneurons were not included in the model, which meant that inhibition received by the pyramidal cells was probably weaker than in the biological situation. Indeed, the pyramidal cells in our model described above (Figures 3 and 4) tended to fire more than they typically do so during theta oscillations *in vivo* (e.g., Soltesz and Deschenes (1993); Robbe et al. (2006)). Is the higher firing frequency of the pyramidal cells related to the weaker inhibition? To answer to latter question, in a subset of the simulations we artificially scaled up inhibition in the model to match the inhibitory synapse numbers on CA1 pyramidal cells that were expected from electron microscopic reconstructions of pyramidal cell dendrites and somata (Megías et al., 2001; Bezaire and Soltesz, 2013). The rationale for scaling up inhibition in this way was that, as described in detail in Bezaire and Soltesz (2013), the estimates of local inhibitory inputs to pyramidal cells were different when based on experimental observations of presynaptic anatomy (local boutons available for synapsing from distinct types of intracellularly filled and reconstructed interneurons) as opposed to postsynaptic anatomy ( inhibitory post-synaptic densities on pyramidal cell dendrites). In simulations with the model containing this rationally scaled up inhibition, only 1% of the pyramidal cells were active, and they fired at a low rate of 1.8 Hz (data not shown), closely resembling the *in vivo* condition (Soltesz and Deschenes, 1993; Robbe et al., 2006). Therefore, the model was capable of reproducing the experimentally observed relatively low firing frequencies for the principal cells during theta oscillations in vivo. However, because the source of the additional inhibition onto CA1 principal cells has not yet been experimentally identified, we used the connectivity estimates as constrained by experimental observations of axonal boutons and lengths in the full scale model (without the scaled-up inhibition) described above (Figures 3 and 4) in the subsequent computational experiments.

### Mechanism of theta generation and phase-preferential firing of interneurons in the full-scale model of the isolated CA1

Next, we examined the onset of the theta rhythm and the firing patterns of the various cell types in the model circuit during theta oscillations (Figure 5 and Table 5). As mentioned above, distinct interneuronal types, defined based on their selective axonal innervation patterns of the postsynaptic domains of pyramidal cells, exhibit characteristic, cell-type-specific preferred phases of firing during theta oscillations *in vivo* (Klausberger et al., 2003, 2004, 2005; Ferraguti et al., 2005; Jinno et al., 2007; Fuentealba et al., 2008; Varga et al., 2012; Lapray et al., 2012; Katona et al., 2014; Varga et al., 2014). Importantly, this fundamental property emerged spontaneously from the full-scale model, without purposeful tuning of parameters except the mean spiking frequency and synaptic strength of the artificial stimulating cells to set the incoming excitation levels from afferents (see Methods for details). As expected, the numerically dominant pyramidal cells, whose intracellular membrane potential oscillations to a large extent generate and underlie the extracellular LFP signal during theta oscillations (Buzsáki et al., 2012), preferentially discharged around the trough 0^*°*^/360^*°*^ of the LFP analog theta rhythm (Figure 5A).

**Figure 5:**
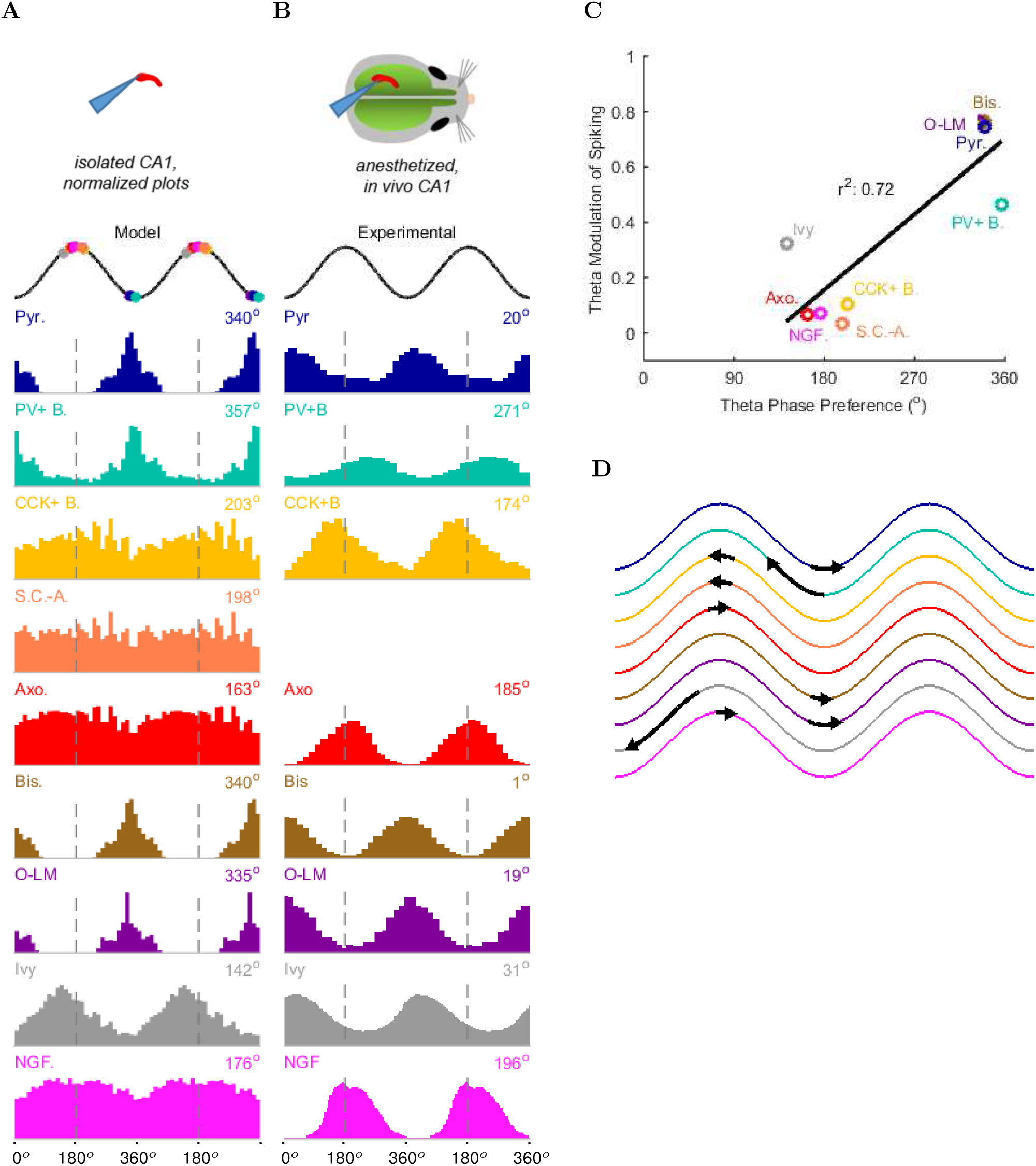
Model and experimental cell theta phases. All model results are based on the spiking of the cells within 100 *µ*m of the reference electrode. (A-B) Firing probability by cell type as a function of theta phase for (A) model and (B) experimental cells under anesthesia (histograms adapted from Klausberger and Somogyi (2008); Fuentealba et al. (2008, 2010) with permission). The model histograms are normalized; see Figure 5 - figure supplement 1 for firing rates. (C) Theta phase preference and theta modulation level were correlated; better modulated cell types spiked closer to the LFP analog trough near the phase preference of pyramidal cells. (D) Theta phase preference plotted on an idealized LFP wave for model data (base of arrow signifies model phase preference and head of arrow shows distance to anesthetized, experimental phase preference). Source Data available at Figure 5 - Source Data.zip.

**Table 5:** Preferred theta firing phases for each model cell type.

| Cell Type | Firing Rate (Hz) | Modulation Level | p | Phase (0°=trough) |
| --- | --- | --- | --- | --- |
| Axo. | 8.9 | 0.07 | 4.58e-130 | 163.4 |
| Bis. | 18.0 | 0.76 | 0.00e+00 | 340.0 |
| CCK+ B. | 54.4 | 0.10 | 0.00e+00 | 202.8 |
| Ivy | 43.3 | 0.33 | 0.00e+00 | 142.1 |
| NGF. | 55.1 | 0.07 | 1.46e-32 | 176.3 |
| O-LM | 17.4 | 0.76 | 0.00e+00 | 334.7 |
| Pyr. | 6.0 | 0.74 | 0.00e+00 | 339.7 |
| PV+ B. | 0.9 | 0.46 | 0.00e+00 | 356.8 |
| S.C.-A. | 5.2 | 0.03 | 1.13e-07 | 197.9 |

Interneurons in the model preferentially fired at specific phases of theta oscillations, depending on the cell type. Their phase preferences fell into two broad categories (Figure 5A). The cells belonging to the first group, including the PV+ basket cells, bistratified cells and O-LM cells, were most likely to fire at the theta trough compared to other theta phases. Since these cells received substantial excitatory inputs from local CA1 pyramidal cells both in the biological state and in the model (Bezaire and Soltesz, 2013), their firing in the isolated CA1 model was probably driven by the pyramidal cell discharges around the theta trough. In contrast, the second group of cells, including the ivy and neurogliaform cells, the CCK+ basket cells and the axo-axonic cells, fired least around the theta trough, leading to an inverted firing probability distribution relative to the first group of interneurons (Figure 5A). Their differing phase preferences were most likely due to a combination of weak or non-existent excitatory inputs from local CA1 pyramidal cells and inhibition from the interneurons that prominently discharged around the theta trough. In general agreement with the first group of cells being strongly and rhythmically driven by the local pyramidal cells, there was a correlation between the phase preference and the strength of modulation (Figure 5C; see Methods), with the cells discharging around the trough all showing strong modulation of firing.

These results were in line with recent data from the isolated CA1 preparation *in vitro* (Ferguson et al., 2015) which showed that cells belonging to the broadly defined SOM+ and PV+ classes (identified using genetic drivers) displayed phase preferences similar to the O-LM, PV+ basket and bistratified cells in our model (note that Ferguson and colleagues used LFP theta recorded in the stratum radiatum as reference, which is approximately 180 degrees out of phase with the pyramidal cell layer theta used in this paper). In addition, the interneuronal phase preferences in the model were also remarkably similar to *in vivo* data from anesthetized animals (Figure 5B; because no data are available on the phase preferential firing of morphologically identified interneurons from the isolated CA1 preparation, comparison is made here with results from anesthetized animals, from which the most complete data sets are available; see also Discussion). specifically, the majority (71%; 5/7) of the interneuron types for which there were experimental data, including the CCK+ basket, axo-axonic, bistratified, O-LM and neurogliaform cells, showed similar preferential maxima in their firing probabilities in the model (Figure 5A) and *in vivo* (Figure 5B). The largest differences between the model and the *in vivo* phase-preferential firing occurred for the PV+ basket cells and the ivy cells, suggesting that during theta oscillations *in vivo* these cells may be strongly driven by CA3 afferents active during the late falling phase of the theta cycle (Colgin and Moser, 2010); note that PV+ cells receive a high number of excitatory inputs on their dendrites compared to other interneuron classes (Gulyas et al., 1999). A comparison of the model and the anesthetized *in vivo* data is illustrated in Figure 5D, where the arrows indicate the shift required for the model phase preferences (Figure 5A) to equal the *in vivo* (Figure 5B) phase preferences; note that the required shifts (arrows) are small for all interneuron types except PV+ basket and ivy cells. A clear majority of the interneuronal types in the model showed phase preferences similar to the *in vivo* condition where rhythmically discharging afferent inputs are present, indicating that theta-preferential discharges are to a large extent determined by the wiring properties of the CA1 circuit itself.

### Perturbation experiments indicate a key role for interneuronal diversity in the emergence of spontaneous theta

Importantly, the ability to generate theta oscillations, phase-locked gamma oscillations, and theta-related phase-preferential firing of distinct interneuronal subtypes was not a universal property of the model. As shown in Figure 6A, our strongly constrained model only exhibited spontaneous theta oscillations at certain levels of afferent excitation. The results described above (Figures 3-5) were obtained with an afferent excitation level of 0.65 Hz (labeled as “Control” in Figure 6A), meaning that each excitatory afferent cell excited the model network with a Poisson-distributed spike train having a Poisson mean interspike interval (ISI) corresponding to a firing rate of 0.65 Hz. When the excitation level decreased below 0.65 Hz, the theta rhythm fell apart, and when the excitation level increased beyond 0.80 Hz, theta power also started to drop significantly as the oscillation frequency rose out of theta range (Figure 6 and Figure 6 - figure supplement 1), evolving into a beta oscillation (Engel and Fries, 2010). These data indicate that while synaptic-cellular organization of the CA1 circuit enables the intrinsic, within-CA1 generation of theta waves, the circuit is predisposed to exhibit theta oscillations only under particular excitatory input conditions. The observation that, under certain conditions the model network can oscillate at frequencies between 12 and 20 Hz, is in agreement with recent experimental findings that rhythmic driving of septal PV+ cells can reliably entrain the hippocampus in a 1:1 ratio up to frequencies of 20 Hz (Dannenberg et al., 2015).

**Figure 6:**
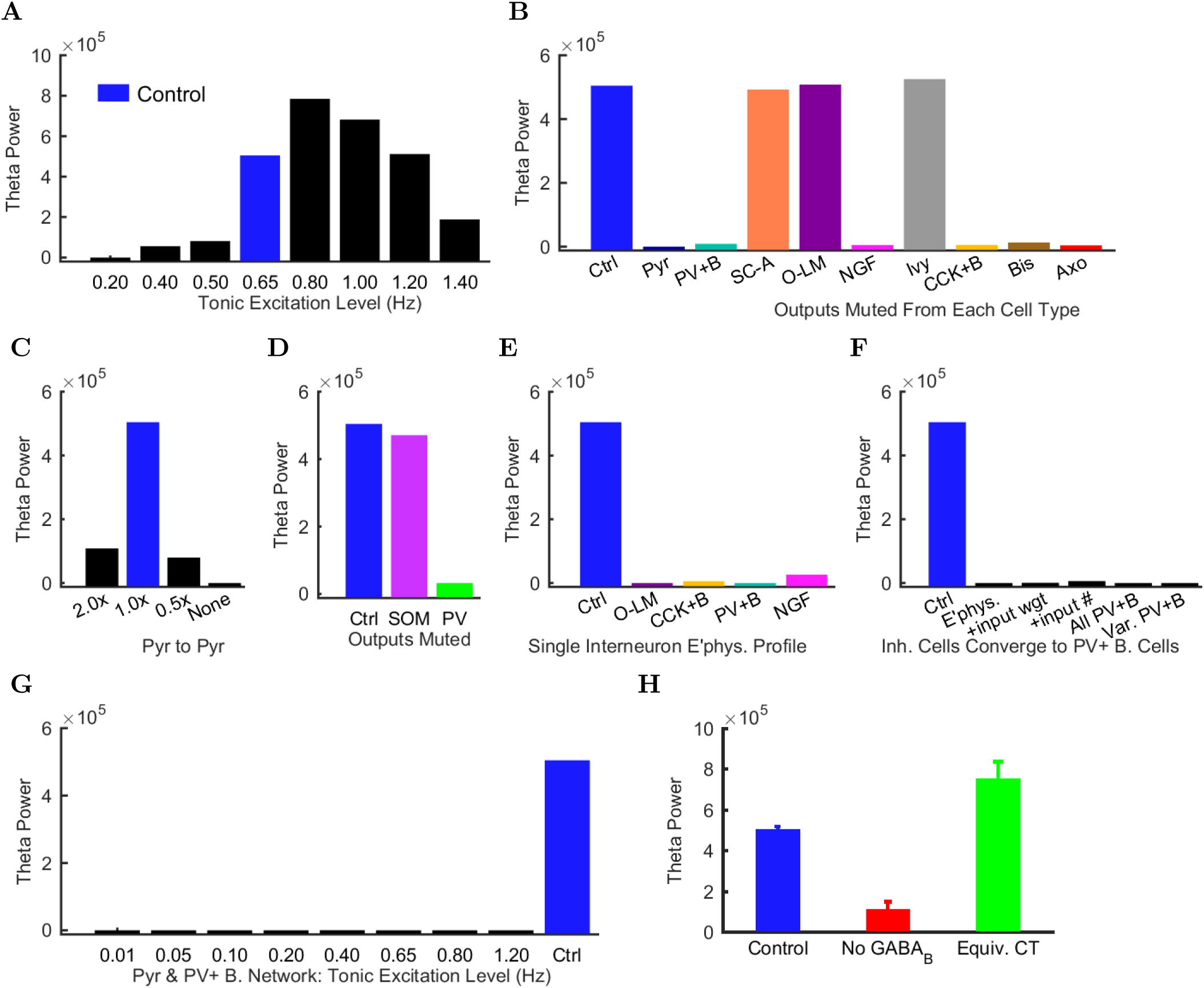
Altered network configurations. Oscillation power (in mV^2^/Hz) of the spike density function (SDF) for pyramidal cells within 100 *µ*m of the reference electrode, at the peak frequency within theta range (5-10 Hz) in altered network configurations. For corresponding peak frequencies, see Figure 6 - figure supplement 1. (A) Theta is present at some excitation levels. (B) Muting each cell type’s output caused a range of effects. (C) The stability and frequency of spontaneous theta in the network was sensitive to the presence and number of recurrent connections between CA1 pyramidal cells. (D) Partially muting the broad classes of PV+ or SOM+ cells by 50% showed that PV+ muting disrupted the network more than SOM+ muting. (E) Theta falls apart when all interneurons are given the same electrophysiological profile, whether it be of a PV+ basket, CCK+ basket, neurogliaform, or O-LM cell. (F) Gradually setting all interneuron properties to those of PV+ basket cells did not restore theta. From left to right: control network; PV+ basket cell electrophysiology; also weights of incoming synapses; also numbers of incoming synapses; then all interneurons being PV+ basket cells (with the addition of the output synapse numbers, weights, and kinetics); then variable RMP (normal disribution with standard deviation of 8 mV). (G) A wide range in excitation was unable to produce theta in the PV+ B. network. (H) Removing the GABA_*B*_ component from the neurogliaform synapses onto other neurogliaform cells and pyramidal cells showed a significant drop in theta power. Massively increasing the weight of the GABA_*A*_ component to produce a similar amount of charge transfer restored theta power (compare the IPSCs corresponding to each condition iny Figure 6 - figure supplement 2). Standard deviations (n=3) shown; significance (p=1.8e-05). Source Data available at Figure 6 - Source Data.zip.

Does the parameter sensitivity of the theta rhythm also apply to recurrent excitation from pyramidal cells and inhibition from CA1 interneurons? In order to answer the latter question, we tested whether the theta rhythm was differentially sensitive to the contribution of each inhibitory cell type (Figure 6B). We characterized the contribution of each local CA1 cell type to the theta rhythm by muting the output of the cell type so that its activity had no effect on the network. First, we studied the role of the recurrent collaterals of pyramidal cells, which contact mostly interneurons and, less frequently, other pyramidal cells (Bezaire and Soltesz, 2013). When we muted all the outputs from pyramidal cells, theta rhythm disappeared (bar labeled “Pyr” in Figure 6B), indicating that the recurrent collaterals of pyramidal cells play a key role in theta oscillations.

Interestingly, muting the relatively rare CA1 pyramidal cell to pyramidal cell excitatory connections alone (each pyramidal cell contacts 197 other pyramidal cells in the CA1; Bezaire and Soltesz (2013)) was sufficient to collapse the theta rhythm (bar labeled “None” in Figure 6C); key roles for inter-pyramidal cell excitatory synapses within CA1 have been suggested for sharp wave ripple oscillations as well (Maier et al., 2011). Furthermore, the parameter-sensitivity of the theta rhythm was also apparent when examining the role of pyramidal cell to pyramidal cell connections, because theta power dramatically decreased when these connections were either increased (doubled) or decreased (halved) from the biologically observed 197 (Figure 6C). Next, we investigated the effects of muting the output from each interneuron type. Silencing the output from any of the fast-spiking, PV family interneurons (PV+ basket, axo-axonic, or bistratified cells), CCK+ basket cells, or neurogliaform cells also strongly reduced theta power in the network (Figure 6B). In contrast, muting other interneuronal types (S.C.-A cells, O-LM cells, or ivy cells) had no effect on this form of theta oscillations generated by the intra-CA1 network (Figure 6B). In additional disinhibition studies simulating optogenetic experimental configurations, partial muting of all PV+ outputs (PV+ basket, bistratified, and axo-axonic cells together) had a larger effect than partial muting of all SOM+ outputs (O-LM and bistratified cells); see Figure 6D. Reassuringly, these results were in overall agreement with experimental data from the isolated CA1 preparation indicating that optogenetic silencing of PV+ cells, but not SOM+ cells such as the O-LM cells, caused a marked reduction in theta oscillations (Amilhon et al., 2015). The differential effects of silencing PV+ versus SOM+ cells could also be obtained in a rationally simplified model called the Network Clamp, where a single pyramidal cell was virtually extracted from the full-scale CA1 network with all of its afferent synapses intact (for further details, see Bezaire et al. (2016)).

Since the diverse sources of inhibition from the various interneuronal types are believed to enable networks to achieve more complex behaviors, including oscillations (Soltesz, 2006; Rotstein et al., 2005; Kepecs and Fishell, 2014), we next tested if reducing the diversity of interneurons in the model would affect its ability to produce spontaneous theta oscillations. Surprisingly, giving all interneurons a single electrophysiological profile appeared to create conditions that were not conducive for the appearance of spontaneous theta oscillations regardless of which interneuronal profile was used (Figure 6E; note that the cells still differed in the strengths, distribution, and identities of their incoming and outgoing connections after this manipulation). To probe this finding further, we focused on PV+ basket cells, which have been implicated in theta generation *in vivo* (Soltesz and Deschenes, 1993; Buzsáki, 2002; Stark et al., 2013; Hu et al., 2014) and exhibited strong theta power in their spiking in the control network model (Figure 4B). We gradually altered (“morphed”) the properties of all other model interneuron types until they became PV+ basket cells, by first converging their electrophysiological profiles, then additionally their synaptic kinetics and incoming synapse weights, then also their incoming synapse numbers, and finally their outgoing synaptic weights and numbers (Figure 6F; Table 7). Theta was not apparent in any intermediate steps nor in the final network where all interneurons had become PV+ basket cells (“All PV+B” in Figure 6F). Furthermore, introduction of cell to cell variability in the resting membrane potential of interneurons in the “All PV+B” configuration at the biologically observed values for PV+ basket cells also failed to restore theta (“Var PV+B” in Figure 6F shows results with standard deviation of (SD) = 8 mV in the resting membrane potential; SD = 5 mV and SD = 2 mV also yielded no theta; biological SD value: approximately 5 mV in Tricoire et al. (2011) and 2 mV in Mercer et al. (2012)). Therefore, although PV basket cells appear to be important for theta-generation both in the biological and the model CA1 network, endowing all interneurons with PV basket cell-like properties does not lead to a network configuration conducive to theta oscillations (Hendrickson et al., 2015).

To rule out the possibility that the lack of theta could be due to an inappropriate excitation level in these reduced diversity configurations, we subjected the “All PV+ B” network to a wide range of incoming excitation levels (Figure 6G). Theta rhythm did not appear at any of these excitation levels. While we could not rule out a hypothetical theta regime somewhere in the parameter space of such low-diversity configurations, any theta solution space would likely be smaller and more elusive than we were able to determine in the control configuration (Figure 6A).

Taken together, these results indicated, for the first time, that interneuronal diversity itself is an important factor in the emergence of spontaneous theta oscillations from the CA1 network.

### Neurogliaform cell signaling and theta generation in the isolated CA1 model

In agreement with previous predictions (Capogna, 2011), the perturbation experiments described above suggested that neurogliaform cells were a necessary component for spontaneous theta to arise in the isolated CA1. We wondered why muting the output from neurogliaform cells, but not the closely related ivy cells, affected theta oscillations (Figure 6B), especially since there were fewer neurogliaform cells than ivy cells, and they were less theta modulated (Figure 5A). These two model interneuron groups mainly differed in that the neurogliaform cells evoked mixed GABA_A,B_ postsynaptic events (Price et al., 2005), whereas the model ivy cells only triggered GABA_A_ IPSPs (in agreement with a lack of evidence for ivy cell-evoked GABA_B_ IPSPs). Could the slow kinetics of GABA_B_ IPSPs contribute to the pacing of the theta oscillations? Indeed, when we selectively removed the GABA_B_ component of all neurogliaform cell outgoing synaptic connections, theta power was strongly reduced (Figure 6H). To test whether the contribution of the GABA_B_ receptors was due to their slow kinetics, we artificially sped up the GABA_B_ IPSPs so that they had GABA_A_ kinetics but conserved their characteristic large charge transfer. This alteration was implemented by scaling up the GABA_A_ synaptic conductance at neurogliaform cell output synapses to achieve a similar total charge transfer as the control GABA_A,B_ mixed synapse (Figure 6 - figure supplement 2). As shown in Figure 6H (green bar), theta activity was restored when the neurogliaform cell output synapses had no slow GABA_B_ component, only a scaled up fast GABA_A_ IPSP with a charge transfer equivalent to the mixed GABA_A,B_ synapses. Therefore, muting the neurogliaform cells strongly disrupted the theta oscillations not because the theta oscillations required the slow kinetics of GABA_B_ IPSPs specifically, but because the slow kinetics enabled a large total charge transfer.

## Discussion

### Emergence of theta oscillations from a biological data-driven, full-scale model of the CA1 network

We produced a biologically detailed, full-scale CA1 network model constrained by extensive experimental data (Bezaire and Soltesz, 2013). When excited with arrhythmic inputs at physiologically relevant levels (see below), the model displayed spontaneous theta (and gamma) oscillations with phase preferential firing across the nine model cell types (pyramidal cells and eight interneuron classes). Consistent with experimental results (Goutagny et al., 2009; Amilhon et al., 2015), these oscillations emerged from the network model without explicit encoding, rhythmic inputs or purposeful tuning of intra-CA1 parameters (all anatomical connectivity parameters were exactly as previously published in Bezaire and Soltesz (2013)). Cell type-specific perturbations of the network showed that each interneuronal type contributed uniquely to the spontaneous theta oscillation, and that the presence of diverse inhibitory dynamics was a necessary condition for sustained theta oscillations. In addition to characterizing roles for specific network components, these model results generally suggest that the presence of diverse interneuronal types and the intrinsic circuitry of the CA1 network are sufficient and necessary to enable the isolated CA1 to oscillate at spontaneous theta rhythms while supporting distinct phase preferences of each class of hippocampal neuron. These abilities may serve to maintain the stability and robustness of the theta oscillation mechanism as it operates *in vivo* in diverse behavioral states. The theta rhythm is thought to be important for organizing disparate memory tasks (Lisman and Idiart, 1995; Hasselmo et al., 2002; Hasselmo, 2005; Lisman and Jensen, 2013; Siegle and Wilson, 2014), and a CA1 network which has evolved a predisposition to oscillate at theta and gamma frequencies may enable more efficient processing of the phasic input it receives *in vivo* (Akam and Kullmann, 2012; Fries, 2015). In turn, phase preferential firing may aid information processing tasks by providing order and allowing multiple channels of information to be processed in parallel (Jensen and Lisman, 2000; Hasselmo et al., 2002; Womelsdorf et al., 2007; Schomburg et al., 2014; Jeewajee et al., 2014; Maris et al., 2016).

Importantly, theta oscillations appeared only within certain levels of excitatory afferent activity, around 0.65 Hz for the average firing rate of the Poisson-distributed spike trains. When the 454,700 stimulating afferents in the model (representing the CA3 and entorhinal synapses; calculated in Bezaire and Soltesz (2013)) are active at a Poisson mean of 0.65 Hz, they generate approximately 37,900 incoming spikes / theta cycle, given a theta frequency of 7.8 Hz (Equation 1).

(1)

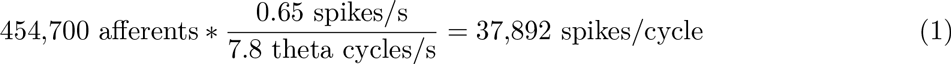

Is the latter number of spikes in the afferents to the CA1 network within a physiologically plausible range? The biological CA1 network receives most of its input from CA3 and entorhinal cortical layer III (ECIII), and it has been estimated that about 4% of CA3 pyramidal cells fire up to four spikes per theta wave (Gasparini and Magee, 2006). We previously estimated 204,700 pyramidal cells in ipsilateral CA3 (Bezaire and Soltesz, 2013), giving an estimated 32,750 spikes from ipsilateral CA3 per theta cycle (Equation 2).

(2)

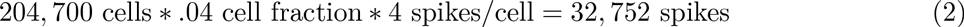

About 250,000 principal cells from ipsilateral ECIII synapse onto the CA1 region (Andersen et al., 2006), and approximately 2% of these cells are active per theta cycle at a low firing rate (Csicsvari et al., 1999; Mizuseki et al., 2009). Therefore, ECIII cells could provide 5,000 input spikes to ipsilateral CA1 (Equation 3).

(3)

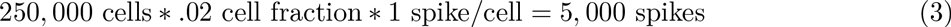

Therefore, about 37,750 spikes per theta cycle arrive from ipsilateral CA3 and entorhinal cortex to the CA1 network *in vivo*, which is reassuringly close to the our modeling results indicating that robust theta emerged when the CA1 network model received approximately 37,900 afferent spikes per theta cycle. Thus, the model has the capacity to process a biologically realistic number of spike inputs per cycle while maintaining the theta rhythm.

Our results obtained using the 0.65 Hz excitation indicated that the CA1 model network exhibited phenomena that corresponded well with experimental results, for example, on the differential roles of PV+ basket cells and OLM cells. In addition, the simulations unexpectedly revealed that interneuronal diversity itself may also be important in theta generation, since conversion of all interneurons into fast spiking PV+ basket cells did not result in a network that was conducive for the emergence of theta, in spite of the key role of the PV+ basket cells in hippocampal oscillations. The modeling results also provided the interesting insight that GABA_B_ receptors may play important roles in slow oscillations such as the theta rhythm not because their slow kinetics pace the oscillations, but because their slow kinetics enable a massive charge transfer. This insight was illuminated by the fact that slow GABA_B_ synapses were not necessary for theta as long as their large charge was carried by the fast GABA_A_ synapses. However, we had to increase the conductance of the GABA_A_ synapse almost 300 times to achieve a similar charge transfer as that conveyed by the GABA_B_ synapse. Such a large conductance is not biologically realistic, indicating that the key role for GABA_B_ synapses may be to allow the temporal distribution of the large synaptic charge transfer. Indeed, the importance of GABA_B_ receptors has also been indicated by a number of recent experimental studies, for example, in the modulation of theta and gamma oscillations (Kohl and Paulsen, 2010), setting of spike timing of neuron types during theta (Kohl and Paulsen, 2010), and playing a role in cortical oscillations and memory processes (Craig and McBain, 2014).

In addition to identifying key roles for certain inhibitory components (PV+ interneurons, neurogliaform cells, GABA_B_, and interneuron diversity), our results also highlighted the importance of the recurrent excitatory collaterals from CA1 pyramidal cells in theta generation in the model of the isolated CA1 network. While it may be expected that isolated theta generation would require local pyramidal cells to provide rhythmic, recurrent excitation to interneurons, our simulations additionally showed that the relatively rare pyramidal cell to pyramidal cell local excitatory connections were also required.

Based on our results, we hypothesize that the inhibitory and excitatory connections within CA1 that were identified to be critical in our perturbation (“muting”) simulations (Figure 6B) interact to generate the theta waves in the model as follows. Pyramidal cells preferentially discharge at the trough of the LFP analog, strongly recruiting especially the PV+ basket and bistratified cells (green and brown raster plots in Figure 3C), which, in turn, cause a silencing of the pyramidal cells (blue raster plot in Figure 3C) for about the first third of the rising half (i.e., from 0° to about 60°) of the LFP analog theta cycle. As the pyramidal cells begin to emerge from this period of strong inhibition, initially only a few, then progressively more and more pyramidal cells reach firing threshold, culminating in the highest firing probability at the theta trough, completing the cycle. The progressive recruitment of pyramidal cells during the theta cycle appears to be paced according to gamma (see blue raster plot in Figure 3C), and it is likely that the intra-CA1 collaterals of the discharging pyramidal cells play key roles in the step-wise (gamma-paced) recruitment of more and more pyramidal cells as the cycle approaches the following trough. The predicted key roles for physiological pyramidal cell to pyramidal cell connections in theta-gamma generation during running may be tested in future experiments.

### Rationale for bases of comparison between modeling results with experimental data

Because our model represented the isolated CA1 network, the modeling results were compared with experimental data from the isolated CA1 preparation when possible. Modeling results for which no corresponding experimental data were available from the isolated CA1 preparation, such as the phase preferential firing of individual interneuron types during theta oscillations, were compared with *in vivo* data from anesthetized animals (Figure 5B). Experimental results from anesthetized animals offered the most complete data set (e.g., no experimental data were available on CCK basket cells and neurogliaform cells from awake animals, see Figure 5 - figure supplement 2). Out of the four interneuronal types for which *in vivo* data were available from both the awake and anesthetized conditions (Figure 5 - figure supplement 2), the phase preference of the axo-axonic cell in the model (163°) was closer to the anesthetized phase (185°) than to the awake phase (251°), whereas the PV+ basket cells in the model displayed phase preferential firing (357°) closer to data reported from awake (289°-310°) than anesthetized animals (234°-271°); the precise reasons underlying these differences are not yet clear. In contrast, bistratified and O-LM cells fired close to the trough in the model, under anesthesia and in awake animals, potentially indicating the primary importance of pyramidal cell inputs in driving these interneurons to fire during theta oscillations under all conditions.

While our model is fundamentally a model of the rat CA1 (e.g., in terms of cell numbers and connectivity; see Table 3 in Bezaire and Soltesz (2013)), some of the electrophysiology data used for constructing the single cell models (Supplementary Material) came from the mouse. In addition, the experimental data on the isolated CA1 preparation were obtained from both rat (Goutagny et al., 2009) and mouse (Amilhon et al., 2015), similar to the experimental results on the phase specific firing *in vivo* (e.g., awake rat: Lapray et al. (2012); awake mouse: Varga et al. (2014)). Because there is no reported evidence for major, systematic differences in key parameters such as the phase specific firing of rat and mouse interneurons in vivo, we did not compare our modeling results with rat and mouse data separately.

A final point concerns the nature of the theta rhythm that emerged in our model. In general, the *in vivo* theta rhythm has been reported to be either atropine resistant or atropine sensitive, where the former is typically associated with walking and may not be dependent on neuromodulatory inputs, while the latter requires intact, rhythmic cholinergic inputs (Kramis et al., 1975). Given that our model did not explicitly represent neuromodulatory inputs, it is likely that the theta that emerged from our model most closely resembled the atropine resistant form. However, it also plausible that both forms of theta benefit from occurring in a network that is predisposed to oscillate at the theta frequency, as the model network results suggested.

### An accessible approach to modeling that balances detail, scale, flexibility and performance

Our results from the strictly data-driven, full-scale CA1 model are consistent with those of earlier models that elegantly demonstrated the basic ingredients capable of producing emergent network oscillations at a range of frequencies in microcircuits and small networks (Rotstein et al., 2005; Siekmeier, 2009; Neymotin et al., 2011b,a; Ferguson et al., 2013). In addition, our modeling approach also provides a full-scale option to advance the recent studies of network activity propagation and information processing during theta (Cutsuridis et al., 2010; Cutsuridis and Hasselmo, 2012; Taxidis et al., 2013; Saudargiene et al., 2015). Here, we demonstrated that emergent theta and gamma oscillations and theta phase preferential firing are possible even as additional interneuron types are incorporated and the network is scaled up to full size with realistic connectivity including 5 billion synapses between the 300,000-plus cells of our network model.

This work is one step in our broader effort to build a 1:1 model of the entire temporal lobe using a hypothesis-driven model development process, where at each stage of model development the models are used to address specific questions. For example, here we employed our newly developed full-scale CA1 model to gain mechanistic insights into the ability of the intra-CA1 circuitry to generate theta oscillations (Goutagny et al., 2009). The current CA1 network model can be developed into a whole hippocampal or temporal lobe model by replacing the designed CA3 and entorhinal cortical afferents with biophysically detailed CA3, ECIII, and septal networks. While we design our model networks with the motivation to answer a particular question, we keep in mind their potential usage for a broad range of questions. Previously, we built a dentate gyrus model to study epileptic network dynamics (Santhakumar et al., 2005; Morgan and Soltesz, 2008) that was then used by several groups to study disparate topics including epilepsy, network mechanisms of inhibition and excitability, simulation optimization, and modeling software (Migliore et al., 2006; Gleeson et al., 2007; Hines et al., 2008a,b; Hines and Carnevale, 2008; Thomas et al., 2009; Winkels et al., 2009; Cutsuridis et al., 2010; Jedlicka et al., 2010a,b; Thomas et al., 2010; Tejada and Roque, 2014). Our previous model has demonstrated how the resource intensive process of designing a detailed, large-scale model is offset by its potential usage in numerous ways by a multitude of groups. On the other hand, future efforts will be needed to continue to incorporate experimental data obtained by the scientific community on additional, not yet represented parameters into the platform offered by our full-scale CA1 network model, e.g., on cell type-specific gap junctions and short-term plasticity, neuromodulators, diversity of pyramidal cells, glial dynamics, cell to cell variability (e.g., Schneider et al. (2014)) and others.

We developed a flexible and biologically relevant model that uses computational resources efficiently, positioning the model to be used by the broader community for many future questions. Importantly, the model can be run on the Neuroscience Gateway, an online portal for accessing supercomputers that does not require technical knowledge of supercomputing (https://www.nsgportal.org/). The model is public, well documented, and also well characterized in experimentally relevant terms (See Supplementary Material and online links given in Methods). In addition, all the model configurations and simulation result data sets used in this work are available online (Bezaire et al., 2015) at (http://doi.org/10.6080/K05H7D60) so the same simulations can easily be repeated with a future, updated model using SimTracker (Bezaire et al., 2016). Mindful that this model could be used by people with a broad range of modeling experience, we have made freely available our custom software SimTracker (http://dx.doi.org/10.1101/081927) that works with the model code to support each step of the modeling process (Bezaire et al., 2016).

## Conclusion and Outlook

As highlighted by the BRAIN Initiative, there is an increasing recognition in neurobiology that we must compile our collective experimental observations of the brain into something more cohesive and synergistic than what is being conveyed in individual research articles if we are to fully benefit from the knowledge that we collectively produce (Ramaswamy et al., 2015; Markram et al., 2015). By assimilating our collective knowledge into something as functional as a model, we can further probe the gaps in our experimental studies, setting goals for future experimental work. On the other hand, as powerful new tools are gathering vast quantities of neuroscience data, the extraction and organization of the data itself are becoming a challenge. At least three large programs are undertaking this challenge: the Hippocampome project (for neuroanatomical and electrophysiological data in the hippocampus of mice; Wheeler et al. (2015)), the Human Brain Project (currently for neuroanatomical and electrophysiological data and models of the rat neocortex, Ramaswamy et al. (2015)), and NeuroElectro (for electrophysiological data from all species and brain areas; Tripathy et al. (2014)). These comprehensive databases create the opportunity to build strongly biology-inspired models of entire networks, with all the cells and synapses explicitly represented. Such models are not subject to the connectivity scaling tradeoff wherein smaller networks have unrealistically low levels of input or unrealistically high connectivity between cells. In addition, such models are usable for investigations into an almost infinite number of questions at any level from ion channels, to synapses, to cell types, to microcircuit contributions. This approach represents a new strategy in computational neuroscience, distinct from and complementary to the use of more focused models whose role is to highlight the potential mechanism of a small number of network components.

The scale, flexibility, and accessibility of our strictly data-driven, full-scale CA1 model should aid the modeling of other large scale, detailed, biologically constrained neural networks. The current CA1 network model produces results in agreement with experimental data, but also extends the results to probe the mechanisms of spontaneous theta generation. It provides specific testable predictions that enable focused design of future experiments, as well as providing an accessible resource for the broader community to explore mechanisms of spontaneous theta and gamma generation. Because the model is available at full scale, it is a relevant resource for exploring the transformation of incoming spatial and contextual information to outgoing mnemonic engrams as part of spatial and memory processing, and other pertinent network dynamics.

## Methods

All results presented in this work were obtained from simulations of computational models. We implemented our CA1 model in parallel NEURON 7.4, a neural network simulator (Carnevale and Hines, 2005). The model simulations were run with a fixed time step between 0.01 and 0.025 ms, for a simulation duration of 2,000 or 4,000 ms (except for Figure 6D where one simulation ran for 1,600 ms). We executed the simulations on several supercomputers, including Blue Waters at the National Center for Supercomputing Applications at University of Illinois, Stampede and Ranger (retired) at the Texas Advanced Computing Center (TACC), Comet and Trestles at the San Diego Supercomputing Center (SDSC), and the High Performance Computing Cluster at the University of California, Irvine. We used our MATLAB-based SimTracker software tool to design, execute, organize, and analyze the simulations (Bezaire et al., 2016).

## Model Development

The CA1 network model included one type of multicompartmental pyramidal cell with realistic morphology and eight types of interneurons with simplified morphology, including PV+ basket cells, CCK+ basket cells, bistratified cells, axo-axonic cells, O-LM cells, Schaffer Collateral-associated cells, neurogliaform cells, and ivy cells.

Model neurons sometimes behave much differently than expected when subjected to current sweep protocols or synaptic inputs that are outside the range of the original protocols used to construct the model. To ensure the model cells exhibited robust biophysical behavior in a wide range of network conditions, we implemented a standard, thorough characterization strategy for each cell type (Supplementary Material).

The behavior of each cell type was characterized using a current injection sweep that matched experimental conditions reported in the literature. Published experimental data was compared side-by-side with model cell simulation results (Supplementary Material). Model cells were connected via NEURON’s double exponential synapse mechanism (Exp2Syn), with each connection comprising an experimentally observed number of synapses (see Table 1).

The connections between cells were determined with the following algorithm, for each postsynaptic and presynaptic cell type combination:

1. Calculate the distances between every presynaptic cell and postsynaptic cell of the respective types;
2. Compute the desired number and distance of connections, as defined by the presynaptic axonal distance distribution and total number of desired connections of this type; the total number of incoming connections expected by each postsynaptic cell type is divided into radial distance bins and distributed among the bins according to the Gaussian axonal bouton distribution of the presynaptic cell;
3. Assign each of the possible connections determined in step 2 (connections within the axonal extent of the presynaptic cell) to their respective distance bins, and randomly select a specific number of connections from each bin (the specific number calculated to follow the axonal bouton distribution).

When determining which cells of the model to connect, we distributed all cells evenly within their respective layers in 3D space and enabled random connectivity for cell connections where the postsynaptic cell body fell within the axonal extent of the presynaptic cell (looking in the XY plane only). Each time a connection was established between two cells, the presynaptic cell innervated the experimentally observed number of synapses on the postsynaptic cell. The synapse locations were randomly chosen from all possible places on the cell where the presynaptic cell type had been experimentally observed to innervate. The random number generator used was NEURON’s nrnRan4int.

## Biological Constraints

The cell number and connectivity parameters were exactly as we reported previously in our in-depth quantitative assessment of anatomical data about the CA1 (Bezaire and Soltesz, 2013). In the latter paper that formed the data-base for the current full-scale model, we combined immunohistochemical data about laminar distribution and coexpression of markers to estimate the number of each interneuron type in CA1. We then extracted from the experimental literature bouton and input synapse counts for each cell type and multiplied these counts by our estimated number of each cell and determined the available input synapses and boutons in each layer of CA1. The number of connections each cell type was likely to make with every other cell type was based on the results of our quantitative assessment. As the quantitative assessment did not make detailed, interneuron type-specific estimates of connections between interneurons, we performed additional calculations to arrive at the numbers of connections between each type of interneuron in our model. Briefly, we determined the number of inhibitory boutons available for synapsing on interneurons within each layer of CA1. Then, we distributed these connections uniformly across the available incoming inhibitory synapses onto interneurons that we had calculated for that layer. We calculated available incoming synapses by using published experimental observations of inhibitory synapse density on interneuron dendrites by cell class and layer in CA1, which we combined with known anatomical data regarding the dendritic lengths of each interneuron type per layer. We therefore made the following assumption: All available incoming inhibitory synapses onto interneurons in a layer have an equal chance of being innervated by the available inhibitory boutons targeting interneurons in that layer. For further details of the exact calculations, please see the Supplementary Material.

The electrophysiology of each cell was tuned using a combination of manual and optimization techniques. We first fit each cell’s resting membrane potential, capacitance, time constant, and input resistance, followed by hyperpolarized properties such as the sag amplitude and time constant, followed by subthreshold depolarized properties such as a transient peak response, and finally active properties such as spike threshold, rheobase, firing rate, action potential width, height, and afterhyperpolarization. For some cells, we employed the Multiple Run Fitter tool within NEURON to simultaneously fit multiple ion channel conductances. The characterization of each cell type, as well as its comparison to experimental data from the same cell type, is included in the the Supplementary Material.

After fitting the cell model properties, we simulated paired recordings to characterize the connections between our model cells. Where experimental data existed for paired recordings, we matched the experimental holding potential and synapse reversal potential, then performed 10 different paired recordings. We characterized the average synapse properties from those 10 runs, including the synaptic amplitude, 10% - 90% rise time, and decay time constant. Finally, we tuned the synaptic weights and time constants to fit our averages to the experimental data.

To determine the synaptic weights and kinetics for those connections that have not yet been experimentally characterized, we used a novel modeling strategy we call Network Clamp, described in Bezaire et al. (2016). As experimental paired recording data were not available to directly constrain the synapse properties, we instead constrained the firing rate of the cell in the context of the *in vivo* network, for which experimental data have been published. We innervated the cell with the connections it was expected to receive *in vivo*, and then sent artificial spike trains through those connections, ensuring that the properties of the spike trains matched the behavior expected from each cell *in vivo* during theta (firing rate, level of theta modulation, preferred theta firing phase). Next, we adjusted the weight of the afferent excitatory synapses onto the cell (starting from experimentally observed values for other connections involving that cell type) until the cell achieved a realistic firing rate similar to had been experimentally observed *in vivo*.

## Stimulation

As none of the model neurons in the CA1 network are spontaneously active, it was necessary to provide excitatory input to them by stimulating their CA3 and entorhinal cortex synapses. Although the model code is structured to allow the addition of detailed CA3 and cortical inputs, the stimulation patterns used in the present study were not representative of the information content thought to be carried via inputs from those areas, because the focus was on the function of the CA1 network in isolation from rhythmic extra-CA1 influences. In accordance with experimental evidence of spontaneous neurotransmitter release (Kavalali, 2015), we modeled the activation of CA3 and entorhinal synapses as independent Poisson stochastic processes. The model neurons were connected to a subset of these afferents, such that they received a constant level of excitatory synaptic input.

We constrained the synapse numbers and positions of the stimulating afferents using anatomical data. To constrain the afferent synapse weights, we used an iterative process to determine the combinations of synaptic weights that enabled most of the interneurons to fire similar to their observed *in vivo* firing rates (Figure 5 - figure supplement 1 and Table 6). First, we used the output of an initial full-scale simulation to run network clamp simulations on a single interneuron type, altering the incoming afferent synapse weights (but not the incoming spike trains) until the interneuron type fired at a reasonable rate. Then, we applied the synaptic weight to the afferent connections onto that interneuron type in the full-scale model. The resulting simulation then led to a new network dynamic as the constrained activity of that interneuron type caused changes in other interneuron activity. We then performed this exercise for each interneuron type as necessary until we achieved a network where all cell types participated without firing at too high of a level. CCK+ cells had a steep response to the weight of the incoming afferent synapses, remaining silent until the weight was increased significantly and then spiking at a high rate, see Figure 5 - figure supplement 1; the particular difficulty in obtaining the in vivo observed firing rate for CCK+ cells in the model may indicate that *in vivo* they may be strongly regulated by extra-CA1 inhibitory inputs (e.g., from the lateral entorhinal cortex; see Basu et al. (2016) that are not included in the isolated CA1 model).

**Table 6:**
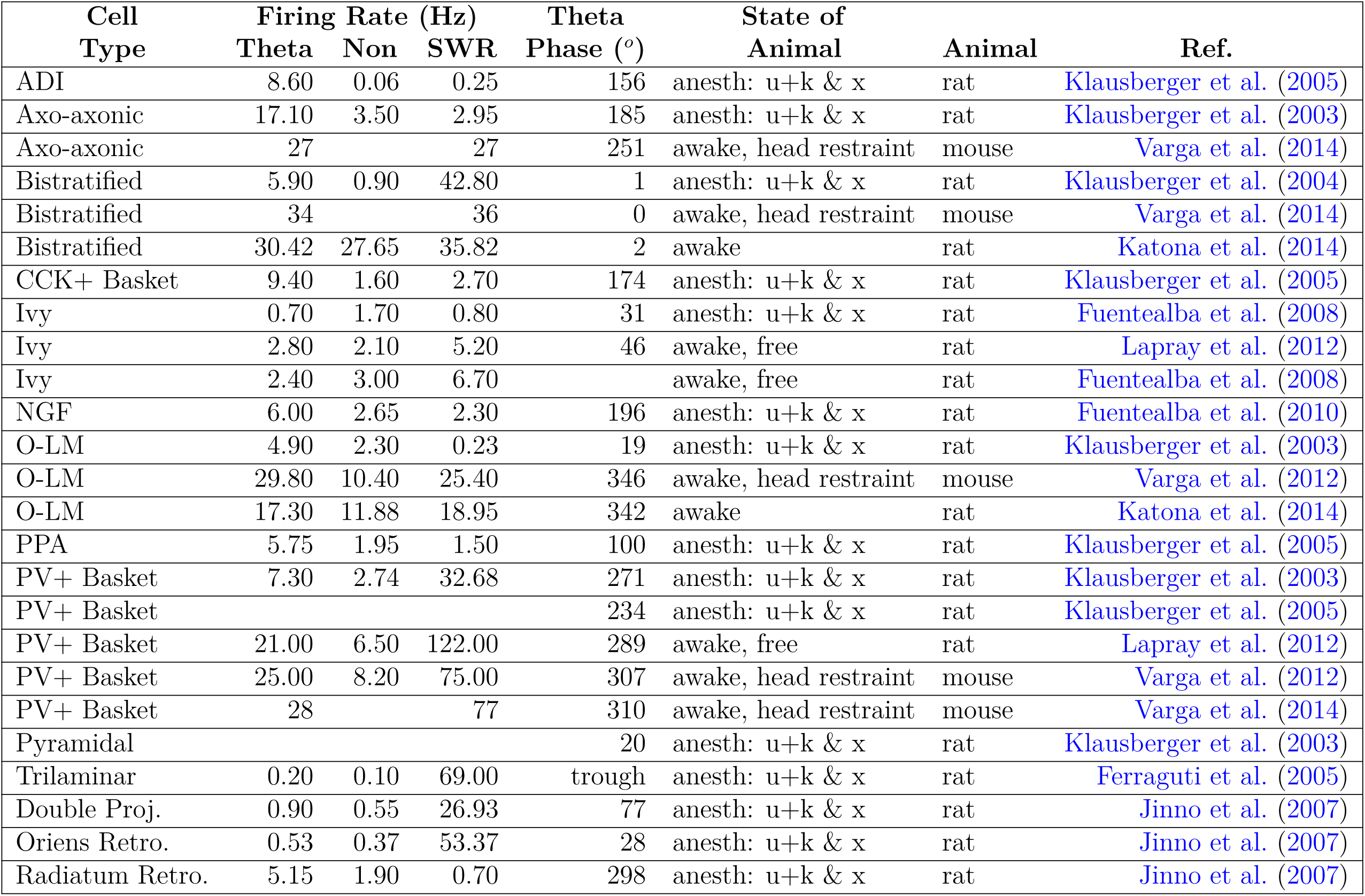
Firing rates and theta phase preferences for various cell types in various conditions. Theta phase is relative to the LFP recorded in the pyramidal layer, where 0° and 360° are at the trough of the oscillation. non: non-theta/non-SWR state. SWR: sharp wave/ripple. u+k & x: urethane + supplemental doses of ketamine and xylazine.

**Table 7:** Peak, theta and gamma frequencies and powers of the pyramidal cell spike density function using Welch’s Periodogram. As in Figure 6 - figure supplement 1, networks where no pyramidal cells spiked - resulting in zero power within the spectral analysis of the pyramidal cell spike density function - have their peak frequencies listed as “n/a” for “not available”.

| Condition | Theta |  | Gamma |  | Overall |  |
| --- | --- | --- | --- | --- | --- | --- |
|  | Frequency | Power | Frequency | Power | Frequency | Power |
| **Tonic Excitation Level (Hz)** |  |  |  |  |  |  |
| 0.20 | n/a | 0.0e+00 | n/a | 0.0e+00 | n/a | 0.0e+00 |
| 0.40 | 5.9 | 5.6e+04 | 25.4 | 4.1e+04 | 13.7 | 6.5e+04 |
| 0.50 | 9.8 | 8.1e+04 | 25.4 | 1.0e+05 | 19.5 | 5.6e+05 |
| 0.65 (Ctrl.) | 7.8 | 5.0e+05 | 25.4 | 2.0e+05 | 7.8 | 5.0e+05 |
| 0.80 | 9.8 | 7.8e+05 | 29.3 | 2.6e+05 | 9.8 | 7.8e+05 |
| 1.00 | 9.8 | 6.8e+05 | 29.3 | 1.4e+05 | 9.8 | 6.8e+05 |
| 1.20 | 9.8 | 5.1e+05 | 33.2 | 1.8e+05 | 11.7 | 8.2e+05 |
| 1.40 | 9.8 | 1.9e+05 | 25.4 | 3.4e+05 | 11.7 | 8.6e+05 |
| **Single Interneuron E'phys. Profile** |  |  |  |  |  |  |
| Ctrl | 7.8 | 5.0e+05 | 25.4 | 2.0e+05 | 7.8 | 5.0e+05 |
| O-LM | n/a | 0.0e+00 | n/a | 0.0e+00 | n/a | 0.0e+00 |
| CCK+B | 9.8 | 5.7e+03 | 62.5 | 6.9e+05 | 62.5 | 6.9e+05 |
| PV+B | n/a | 0.0e+00 | n/a | 0.0e+00 | n/a | 0.0e+00 |
| NGF | 5.9 | 2.6e+04 | 39.1 | 2.4e+06 | 39.1 | 2.4e+06 |
| **Inh. Cells Converge to PV+ B. Cells** |  |  |  |  |  |  |
| Ctrl | 7.8 | 5.0e+05 | 25.4 | 2.0e+05 | 7.8 | 5.0e+05 |
| E'phys. | n/a | 0.0e+00 | n/a | 0.0e+00 | n/a | 0.0e+00 |
| +input wgt | 7.8 | 6.8e+02 | 44.9 | 1.6e+06 | 21.5 | 3.4e+06 |
| +input # | 9.8 | 6.1e+03 | 31.3 | 1.1e+06 | 15.6 | 2.0e+06 |
| All PV+B | n/a | 0.0e+00 | n/a | 0.0e+00 | n/a | 0.0e+00 |
| Var. PV+B | n/a | 0.0e+00 | n/a | 0.0e+00 | n/a | 0.0e+00 |
| **Outputs Muted** |  |  |  |  |  |  |
| Ctrl | 7.8 | 5.0e+05 | 25.4 | 2.0e+05 | 7.8 | 5.0e+05 |
| SOM | 7.8 | 4.7e+05 | 27.3 | 1.4e+05 | 7.8 | 4.7e+05 |
| PV | 9.8 | 3.2e+04 | 27.3 | 8.1e+05 | 13.7 | 1.5e+06 |
| **Pyr to Pyr** |  |  |  |  |  |  |
| 2.0x | 9.8 | 1.1e+05 | 25.4 | 7.3e+05 | 13.7 | 1.0e+06 |
| 1.0x (Ctrl.) | 7.8 | 5.0e+05 | 25.4 | 2.0e+05 | 7.8 | 5.0e+05 |
| 0.5x | 7.8 | 8.0e+04 | 29.3 | 2.2e+05 | 29.3 | 2.2e+05 |
| None | 9.8 | 1.1e+00 | 70.3 | 3.7e+01 | 70.3 | 3.7e+01 |
| **Outputs Muted From Each Cell Type** |  |  |  |  |  |  |
| Ctrl | 7.8 | 5.0e+05 | 25.4 | 2.0e+05 | 7.8 | 5.0e+05 |
| Pyr | 7.8 | 1.1e+00 | 70.3 | 3.8e+01 | 70.3 | 3.8e+01 |
| PV+B | 9.8 | 8.8e+03 | 29.3 | 1.9e+06 | 29.3 | 1.9e+06 |
| SC-A | 9.8 | 4.9e+05 | 27.3 | 1.8e+05 | 9.8 | 4.9e+05 |
| O-LM | 7.8 | 5.1e+05 | 25.4 | 8.3e+04 | 7.8 | 5.1e+05 |
| NGF | 9.8 | 5.2e+03 | 27.3 | 9.1e+05 | 13.7 | 1.6e+06 |
| Ivy | 7.8 | 5.3e+05 | 25.4 | 2.0e+05 | 7.8 | 5.3e+05 |
| CCK+B | 5.9 | 5.5e+03 | 25.4 | 3.3e+03 | 3.9 | 5.7e+03 |
| Bis | 5.9 | 1.3e+04 | 29.3 | 1.7e+06 | 29.3 | 1.7e+06 |
| Axo | 7.8 | 4.0e+03 | 33.2 | 1.2e+06 | 15.6 | 1.9e+06 |
| **Pyr & PV+ B. Network: Tonic Excitation Level (Hz)** |  |  |  |  |  |  |
| 0.01 | n/a | 0.0e+00 | n/a | 0.0e+00 | n/a | 0.0e+00 |
| 0.05 | n/a | 0.0e+00 | n/a | 0.0e+00 | n/a | 0.0e+00 |
| 0.10 | n/a | 0.0e+00 | n/a | 0.0e+00 | n/a | 0.0e+00 |
| 0.20 | n/a | 0.0e+00 | n/a | 0.0e+00 | n/a | 0.0e+00 |
| 0.40 | 5.9 | 2.3e+02 | 25.4 | 1.2e+02 | 3.9 | 2.4e+02 |
| 0.65 | n/a | 0.0e+00 | n/a | 0.0e+00 | n/a | 0.0e+00 |
| 0.80 | n/a | 0.0e+00 | n/a | 0.0e+00 | n/a | 0.0e+00 |
| 1.20 | n/a | 0.0e+00 | n/a | 0.0e+00 | n/a | 0.0e+00 |
| Ctrl | 7.8 | 5.0e+05 | 25.4 | 2.0e+05 | 7.8 | 5.0e+05 |

## Analysis of Simulation Results

We analyzed the results of each simulation with standard neural data analysis methods provided by our SimTracker software (Bezaire et al., 2016), including the spike density function (SDF) of all pyramidal neuron spikes (Szűcs, 1998), the periodogram of the SDF, and the spectrogram of the LFP analog. We determined the dominant theta and gamma frequencies for the network as the peak in the power spectral density estimate obtained by the spectrogram, and confirmed that those peaks are identical for the SDF and the LFP analog. After finding a dominant theta or gamma frequency, we then analyzed the level of modulation and preferred firing phase for each cell type. Finally, we calculated the firing rate of each cell type.

## LFP analog

We calculated an approximation of the LFP generated by the model neurons based on the method described by Schomburg et al. (2012). For each pyramidal cell within 100 *µ*m of a reference electrode location in stratum pyramidale (coordinates = longitudinal: 200 *µ*m; transverse: 500 *µ*m; height from base of stratum oriens: 120 *µ*m), the contribution to extracellular potential at each point along the dendritic and axonal morphology was recorded using NEURON’s extracellular mechanism and scaled in inverse proportion to the distance from the electrode. In order to reduce the computational load of the simulation, 10% of the pyramidal cells outside the 100 *µ*m radius were randomly selected; their distance-scaled extracellular potentials were scaled up by a factor of 10 and then added to the contributions of the inner cells. We performed reference simulations and LFP analog calculations with the inner radius set to 200 *µ*m and 500 *µ*m and obtained results identical with those in Figures 3 and 4 (where an inner radius of 100 *µ*m was used), except for negligible increases in the theta oscillation power found in the LFP analog spectrogram.

## Spike Density Function

We calculated the spike density function (SDF) of all pyramidal cell spikes using a Gaussian kernel with a window of 3 ms and a bin size of 1 ms (Szűcs, 1998). To see how a cell’s spiking activity is related to its SDF, see Figure 4 - figure supplement 1.

## Oscillations

To quantify the frequency and power of the oscillations of the network, we computed a one-sided Welch’s Periodogram of the SDF (Colgin et al., 2009) using a Hamming window with 50% overlap. To characterize the stability of the theta oscillation, we ran the control network for 4 seconds and then computed the spectrogram of the SDF and of the LFP analog using an analysis script from Goutagny et al. (2009) based on the mtspecgramc function from the Chronux toolbox (http://chronux.org/).

## Spike Phases and Theta Modulation

We calculated the preferred firing theta phases of each cell, using all the spikes of that cell type that occurred after the first 50 ms of the simulation, relative to the filtered LFP analog. The spike times were converted to theta phases, relative to the troughs of the LFP analog theta cycle in which they fired. We then subjected the spike phases to a Rayleigh test to determine the level of theta modulation of the firing of each cell type (Varga et al., 2014).

## Firing Rates

The firing rates of the cells were calculated by cropping the first 50 ms of the simulation to remove the initial effects, and then dividing the resulting number of spikes of each cell type by the total number of cells of that type and the duration of the simulation. An alternate average firing rate was calculated by dividing by the number of active cells of that type rather than all of the cells of that type, which gave the average firing rate over all firing cells instead, to better compare with experimentally observed firing rate averages.

## Statistical Comparison of Theta Power

For the GABA_B_-related simulations, we ran three of each condition and then performed an ANOVA to test for significance in the difference of theta power among the conditions.

## Cross correlation of theta and gamma

To investigate whether a relationship existed between the simultaneous theta and gamma oscillations found in the LFP analog of our control simulation, we filtered the LFP analog signal within the theta range (5-10 Hz) and the gamma range (25-80 Hz). We applied a Hilbert transform to each filtered signal and then compared the phase of the theta-filtered signal with the envelope of the gamma-filtered signal to determine the extent to which theta could modulate the gamma oscillation.

## Accessibility

Our model code is available online at ModelDB (code version used to produce results in this work: https://senselab.med.yale.edu/ModelDB/showModel.cshtml?model=187604) and Open Source Brain (most recent code version: http://opensourcebrain.org/projects/nc_ca1). Open Source Brain provides tools for users to characterize and inspect model components. The model is also characterized online at http://mariannebezaire.com/models/ca1, along with a graphical explanation of our quantitative assessment used to constrain the model connectivity Bezaire and Soltesz (2013), as well as links to our model code and model results, and detailed instruction manuals for our NEURON code and SimTracker tool (Bezaire et al., 2016).

For those who wish to view and analyze our simulation results without rerunning the simulation, our simulation results are available on CRCNS.org at http://doi.org/10.6080/K05H7D60 and can be freely accessed after obtaining a free account (Bezaire et al., 2015). Our analyses of these data can be recreated using our publically available SimTracker tool.

Our custom software tool, SimTracker is further discussed in our companion paper, Bezaire et al. (2016). SimTracker is freely available online at http://mariannebezaire.com/simtracker/ and is also listed in SimToolDB at https://senselab.med.yale.edu/SimToolDB/showTool.cshtml?Tool=153281. The tool is offered both as a stand-alone, compiled version for those without access to MATLAB (for Windows, Mac OS X, and Linux operating sytems), and as a collection of MATLAB scripts for those with MATLAB access.

## Acknowledgements

Immeasurable support was provided by NEURON developers Michael Hines and Ted Carnevale under NIH NINDS grant R01-NS11613 (to M.H.) and NSF grant 1458495 (to T.C.). This work used the Extreme Science and Engineering Discovery Environment (XSEDE), which is supported by National Science Foundation grant number ACI-1053575; the project was supported by XSEDE Research Allocation grant TG-IBN140007 (to I.S., M.B. and I.R.) and XSEDE Startup Allocations to
I.S. (TG-IBN130022) and M.B. (TG-IBN100011) and via the Neuroscience Gateway with the support of NSF grants 1458840 and 1146949 (to Majumdar et al.). The authors acknowledge the Texas Advanced Computing Center (TACC) at The University of Texas at Austin for providing high performance computing resources that have contributed to the research results reported within this paper (http://www.tacc.utexas.edu). Additionally, this research is part of the Blue Waters sustained-petascale computing project, which is supported by the National Science Foundation (awards OCI-0725070 and ACI-1238993) and the state of Illinois. Blue Waters is a joint effort of the University of Illinois at Urbana-Champaign and its National Center for Supercomputing Applications (NCSA). Parallel supercomputers used in this work include: Blue Waters, owned by the University of Illinois and NCSA; Stampede and the retired Ranger, owned by the University of Texas’ Texas Advanced Computing Center (TACC); Trestles and Comet, owned by the San Diego Supercomputing Center; University of California at Irvine’s High Performance Computer and the retired Broadcom Distributed Unified Cluster.

We would like to thank the University of Texas’ Texas Advanced Computing Center team, the San Diego Supercomputing Center and Neuroscience Gateway teams (especially Glenn Lockwood, Amitava Majumdar, Subhashini Sivagnanam, Mahidhar Tatineni, and Kenneth Yoshimoto), and UC Irvine’s HPC team (especially Joseph Farran and Harry Mangalam) for their excellent technical support throughout this work. We would also like to thank Padraig Gleeson, Andras Ecker, Tom Morse, and Jeff Teeters for assistance making our code and model results public, and Jesse Jackson and Sylvain Williams for the use of their spectrogram analysis script.

## Figure Supplements

**Figure 1 - figure supplement 1:**
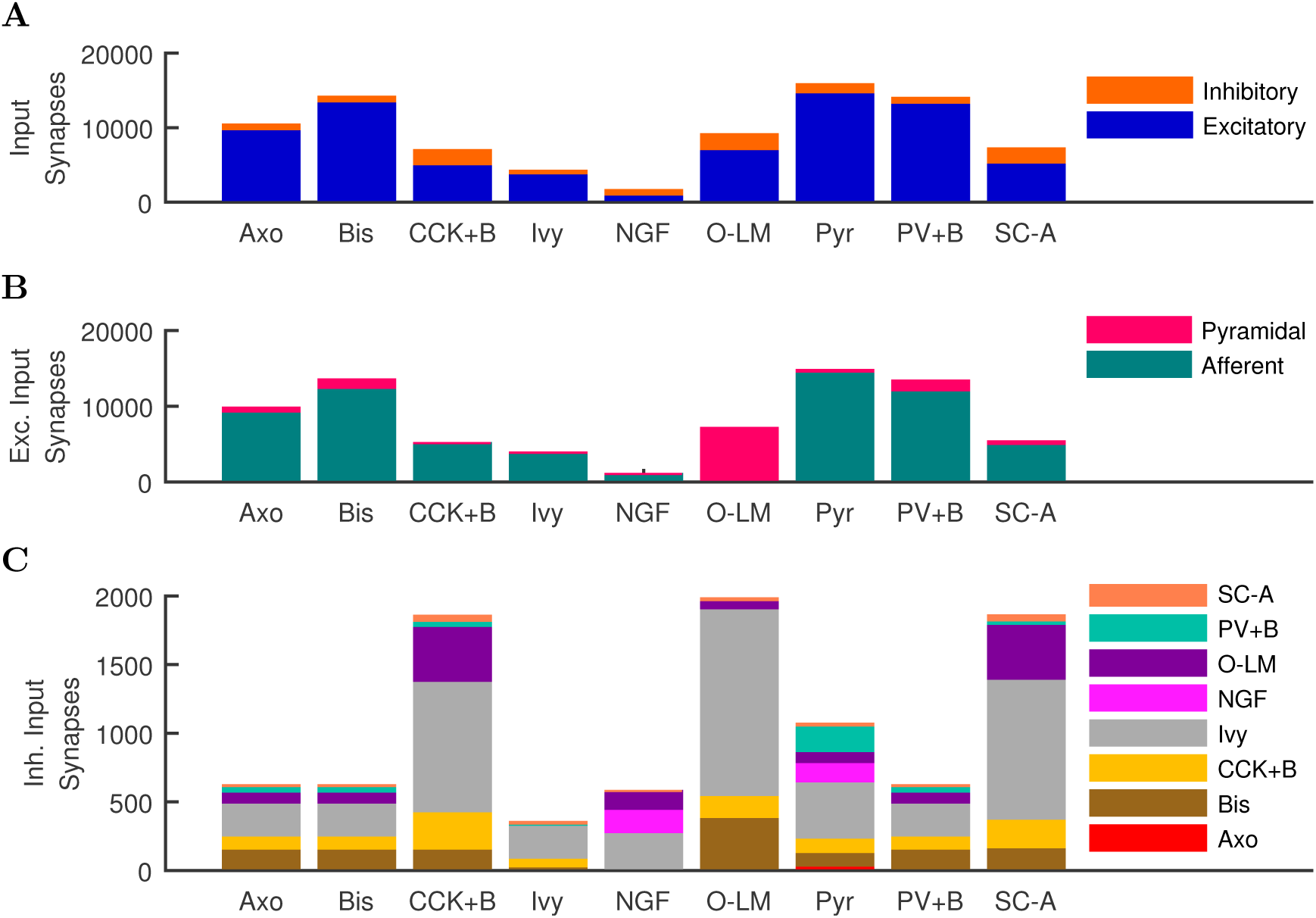
Quantitative Network Connectivity. The average number of incoming synapses per postsynaptic cell of the given type are shown for (A) all inputs to the cells, (B) all excitatory inputs to the cells and (C) all inhibitory inputs to the cells.

**Figure 1 - figure supplement 2:**
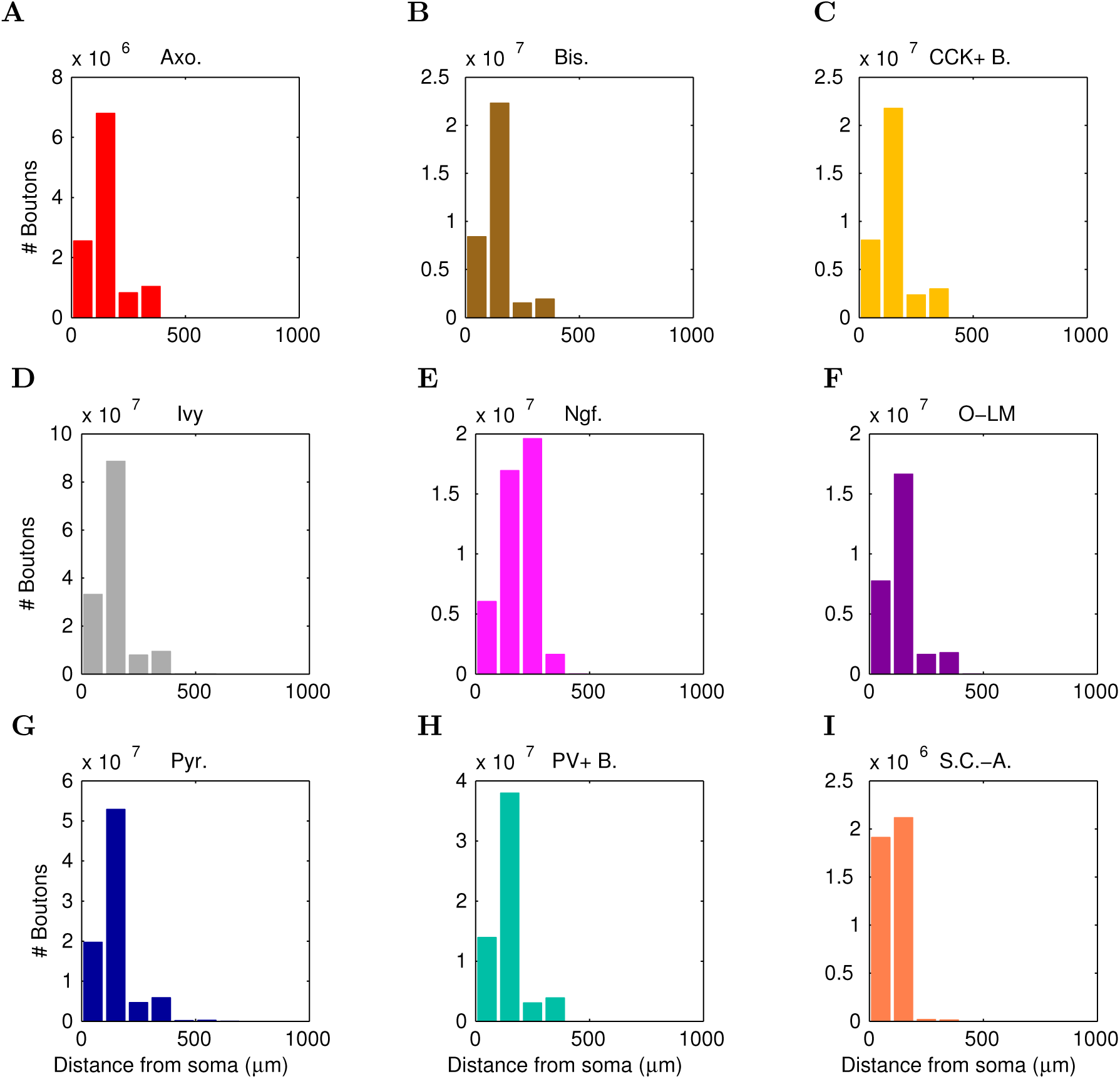
Anatomically constrained connectivity. The axonal distributions are shown per presynaptic cell type. The distribution of boutons is plotted as a function of distance from the presynaptic cell’s soma. Boutons connecting to all possible types of postsynaptic cells are included in the plot. The colors correspond to each presynaptic cell type using the same color code as previous figures.

**Figure 4 - figure supplement 1:**
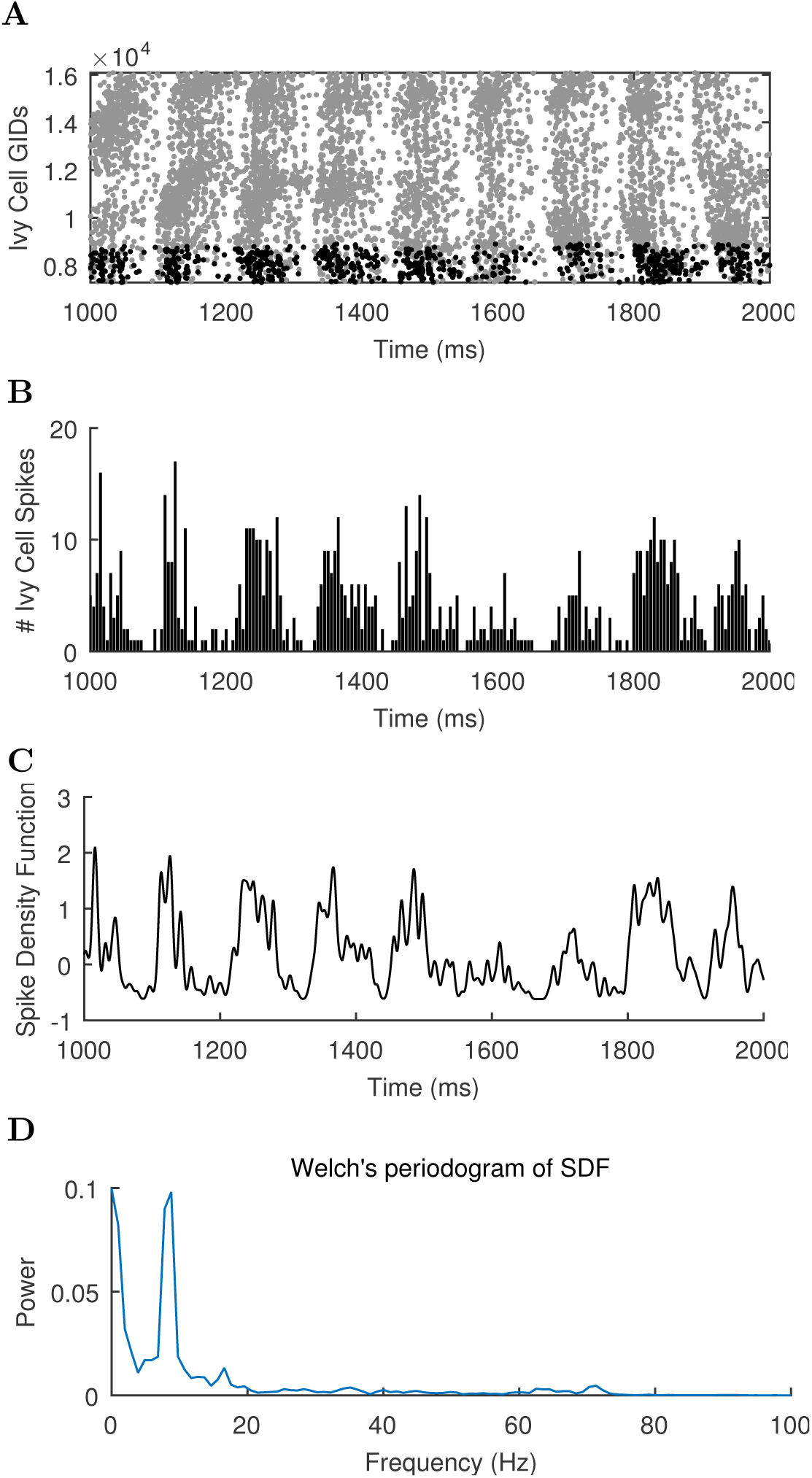
Different views of cell activity. Several ways of characterizing model cell activity per cell type are shown using the spikes from the ivy cells as an example. (A) The spike times of each ivy cell are plotted as a function of time and ivy cell number. A subset of ivy cells positioned within 100 *µ*m of the reference electrode location (whose spikes are shown in black) are then carried forward in the remaining calculations. (B) The spikes of the local ivy cells are binned into 1 ms windows to give spike counts per window. (C) A continuous representation of the ivy cell spikes as a function of time is given in the spike density function (SDF) computed from the ivy cell spike times. (D) A Welch’s Periodogram is computed, which summarizes the power of each oscillation frequency in the ivy cell SDF Although only a part of the simulation is shown, the full simulation length (except the first 50 ms) was used in the spectral analysis.

**Figure 5 - figure supplement 1:**
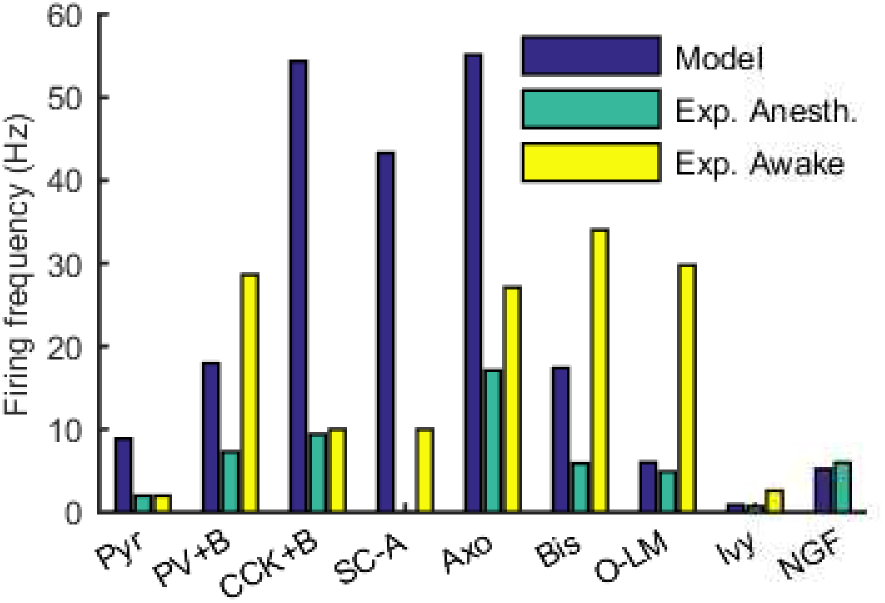
Firing rates of model and experimental cells of each type. For experimental cells, firing rates in both the anesthetized and awake states were included where available. See Table 6 for sources of experimental data.

**Figure 5 - figure supplement 2:**
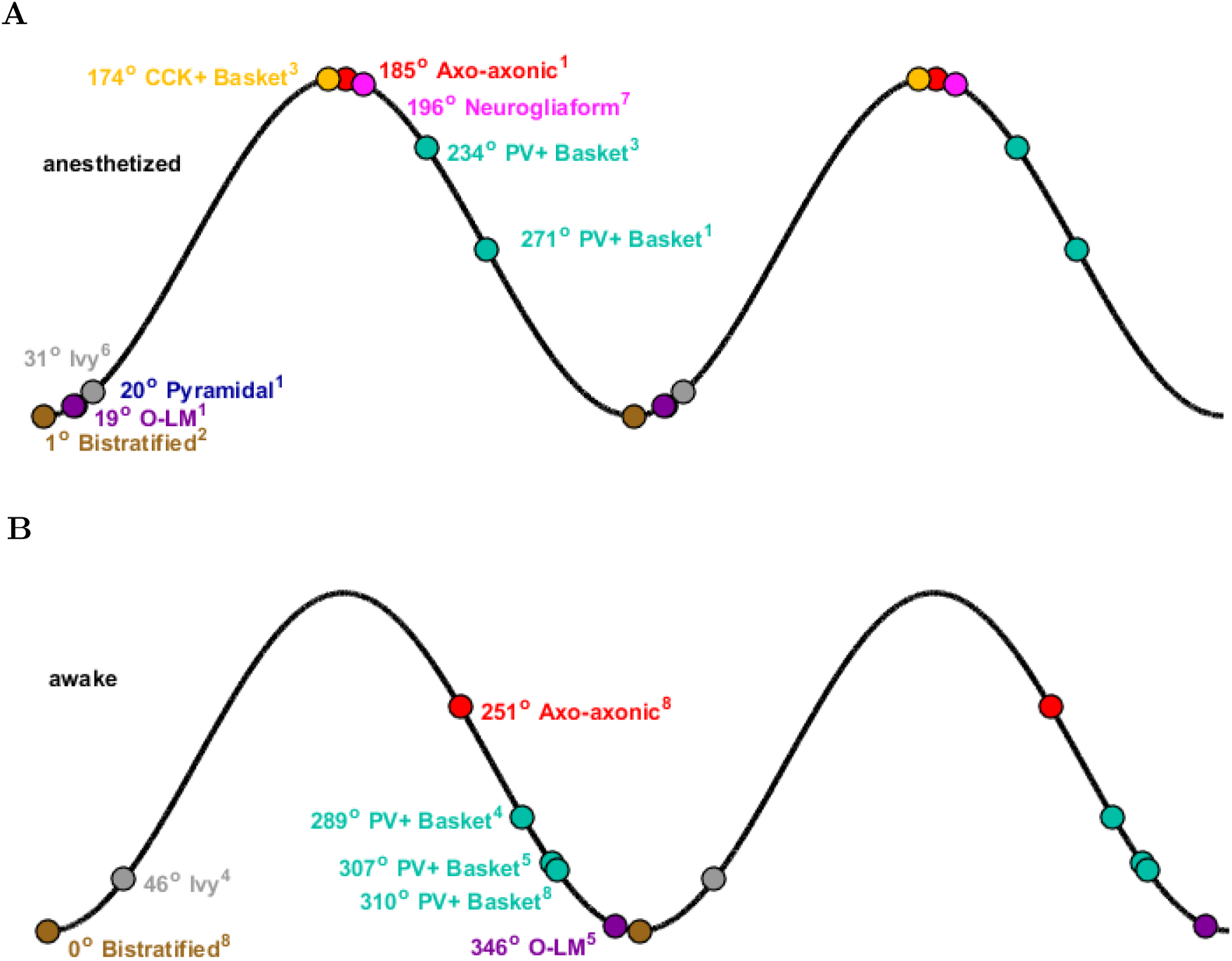
Theta phase-specific firing preferences of various biological hippocampal cell types as reported in the literature. The trough of the pyramidal-layer LFP is designated as 0*°*/360*°* and the peak as 180*°*. There is variation in phase preference for given cell types as a function of experimental preparation. Shown are (A) anesthetized and (B) awake experimental conditions. Reference subscripts correspond to: 1: Klausberger et al. (2003), 2: Klausberger et al. (2004), 3: Klausberger et al. (2005), 4: Lapray et al. (2012), 5: Varga et al. (2012), 6: Fuentealba et al. (2008), 7: Fuentealba et al. (2010), 8: Varga et al. (2014). See Table 6 for further details.

**Figure 6 - figure supplement 1:**
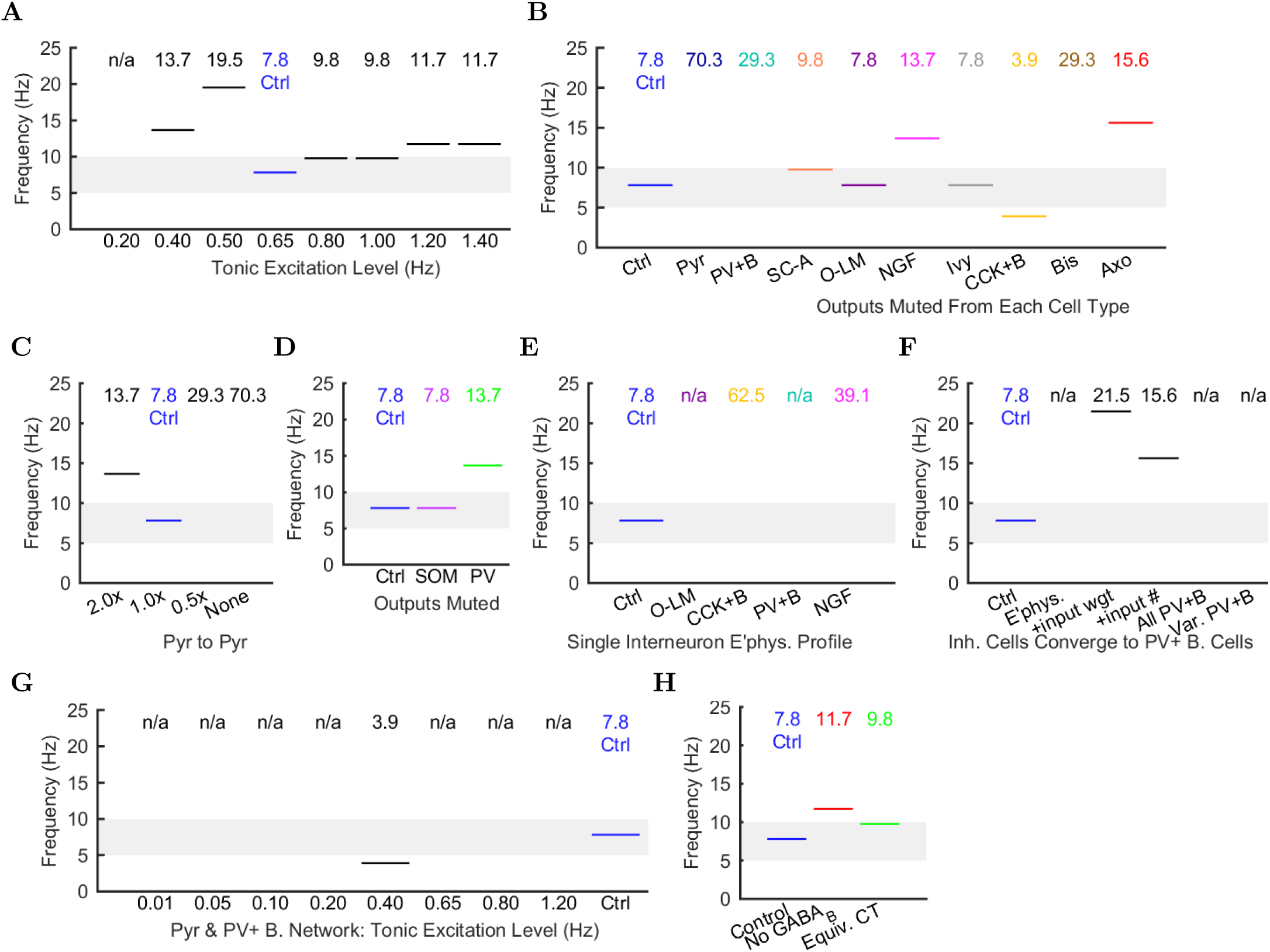
Peak Frequencies of Oscillations in Altered Networks. Peak theta frequency (within 5-10 Hz) of the spike density function (SDF) for all pyramidal cells within 100 *µ*m of the reference electrode in each altered network configuration. For networks where no pyramidal cells spiked, resulting in zero power within the spectral analysis of the pyramidal cell spike density function, their peak frequencies are listed as “not available” or “n/a”. (A) Spontaneous theta oscillation accelerated out of theta range with more excitation. (B) Muting each cell type shifted the oscillation out of range (neurogliaform, CCK+ basket, and axo-axonic cells), disrupted theta but not gamma (not shown; pyramidal, PV+ basket, and bistratified cells), or had little effect (S.C.-A., O-LM, and ivy cells). (C) Doubling the connections between CA1 pyramidal cells increased the theta frequency, while networks with half the number or no recurrent collaterals lost the slow oscillation but kept gamma. (D) Removing 50% of PV+ cell inhibition (PV+ basket, bistratified, and axo-axonic cells) or 50% of SOM+ cell inhibition (bistratified or O-LM cells) shifted the oscillation out of theta range or lost the slow oscillation entirely but kept gamma. (E) Peak oscillation shifted out of theta range when all interneurons had the same electrophysiological profile, regardless of the profile used. (F) Converging all properties to PV+ basket cells, gamma was restored (not shown) but not theta (left to right: control; network with 1: diverse interneurons with same electrophysiology; 2: also with same weights of incoming synapses; 3: also with same numbers of incoming synapses; 4: complete conversion to PV+ basket cells; 5: added variability in resting membrane potential (normal distribution with st. dev. = 8 mV)). (G) In the all-PV+ basket cell network, a wide range of excitation levels could not produce a spontaneous theta rhythm. (H) Removing GABA_B_ increased the oscillation frequency.

**Figure 6 - figure supplement 2:**
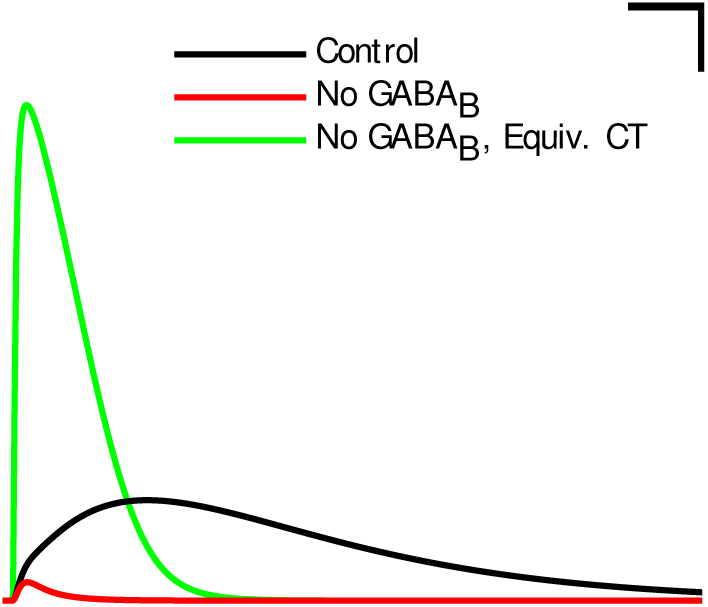
**IPSCs from the neurogliaform to pyramidal cell synapse corresponding to the different conditions in Figure 6H.** These traces are from pyramidal cells clamped at -50 mV during a paired recording from a presynaptic neurogliaform cell with a GABA_A_ reversal potential of -60 mV and a GABA_B_ reversal potential of -90 mV. The currents shown are averages from 10 recordings. Scale bar = 100 ms and 5 pA.

## Source Data Files

### Figure 2 - Source Data

Included are data for voltage and current clamp data for ion channel, single cell, and synaptic characterizations. For the four interneuron types with current injection sweeps displayed, a separate file is provided for each injection in the sweep, with the naming convention trace_[cell type]_[current injection level].dat, where the current injection level is given in pA. For all interneuron types, the current injection sweep data has also been gathered into an AxoClamp ATF (tab-delimited) style of file, to allow for (re)calculation of cell properties according to the processes used for calculations from biological recordings. These files follow the naming convention of [celltype].atf.

The ion channel characterized in this figure was an Na_*v*_ channel, inserted into a single compartment cell of diameter and length 16.8 microns (a soma) with a density such that the maximum, macroscopic conductance was .001 *µ*S/cm^2^. The reversal potential of the channel was +55 mV and the settings during the characterization protocol were: temperature=34 degrees Celsius, axial resistance = 210 ohm*cm, [Ca2+]_internal_ = 5.0000e-06 mM, specific membrane capacitance = 1 *µ*F/cm^2^. For activation steps, the cell was held at -120 mV and then stepped to potential levels ranging from -60 mV to +80 mV. For inactivation steps, the cell was held at various potential levels ranging from -120 mV to +40 mV for 500 ms and then stepped to +20 mV. Each current injection step is recorded in a separate file, with activation step files following the name convention of stepto_[stepped-to potential in mV].dat and inactivation step files following the name convention of hold_[held-at potential in mV].dat.

For the synaptic responses, the postsynaptic cell was voltage-clamped at -50 mV and the reversal potential of the synapse was kept at its natural (as defined in the network model code) potential. A spike was triggered in the presynaptic cell and the current response was measured in the postsynaptic cell at the soma. This recording was repeated 10 times, with a randomly chosen connection location each time, and the response was then averaged. In all paired recordings with the pyramidal cell as postsynaptic cell, the sodium channels were blocked to prevent a suprathreshold response. The file convention for paired recordings is [presynaptic cell].[postsynaptic cell].[Recording #].dat

- stepto_[stepped-to potential in mV].dat
- hold_[held-at potential in mV].dat
- trace_[cell type]_[current injection level].dat
- [celltype].atf

### Figure 3 - Source Data

This zip file contains 3 files. First, it includes the LFP.dat file which contains the raw, theta-filtered, and gamma-filtered LFP analog traces (the raw local field potential (LFP) analog was calculated from the network activity as detailed in the Methods section). Second, the zip file contains Membrane_Potentials.txt, which includes the full duration, intracellular somatic membrane potential recordings from the specific cells shown in Figure 3. Third, it includes the SpikeRasterLocal.dat file which includes the spike times for the length of the entire simulation, from the specific cells displayed in raster shown in Figure 3. The spike times of every single cell in the network are available in the CRCNS repository. Note that the displayed spike raster in Figure 3 has been downsampled in such a way as to preserve its visual appearance while reducing the image size and load time.

- FilteredLFP.dat
- Membrane_Potentials.txt
- SpikeRasterLocal.dat

### Figure 4 - Source Data

This zip file contains the All_SDF_Ctrl_Condition.txt file. The file includes the Spike Density Functions (SDFs) of each cell type in the control network. The calculation of the Spike Density Function is detailed in the Methods section. The power spectra of the SDFs shown in Figure 4 were obtained via a one-sided periodogram using Welch’s method where segments have a 50% overlap with a Hamming Window. The spectra for each cell type was normalized to itself so that each cell type could use the full range of colors in the colorbar to show the shape of its spectra, despite different absolute peak powers for different cell types. Note that the specrogram was computed from the raw LFP analog trace (included in the source data of Figure 3) using the method detailed in Goutagny et al (2009). The cross-frequency coupling was illustrated by performing a Hilbert transform on the raw LFP analog trace to extract the theta phase and gamma envelope as a function of time.

- LFP.dat
- All_SDF_Ctrl_Condition.txt

### Figure 5 - Source Data

This zip file contains a celltype.txt file that summarizes information about each cell type in the model, and it contains one file per each cell type, with the name convention Spike_Phase_Time_[cell type].txt. The Spike_Phase_Time_* files list, for each cell type, the spike times from all of the cells of that type, and calculated theta phases (relative to the theta-filtered LFP analog) of each spike. From the list of spike phases per cell type, the preferred theta phase, level of theta modulation, and statistical significance were calculated for each cell type using a Rayleigh test for circular data (Varga et al, 2014). The average firing rates of each cell type were obtained by dividing the total number of spikes by the length of the simulation (4000 ms) and by the number of cells of each type (listed in the celltype.txt file).

- Spike_Phase_Time_*cell.txt
- celltype.txt

### Figure 6 - Source Data

This zip file contains two files. First, it includes a tab-delimited text file called Pyramidal_SDF_All_Conditions.txt, which contains the full length Spike Density Function computed at a resolution of 1000 Hz from the spikes of all pyramidal cells within the local range of the electrode point in the model network, for each network condition studied in Figure 6. Second, it contains a file called Mapping_Network_Condition.txt that maps the names of the simulations (used in the header of Pyramidal_SDF_All_Conditions.txt) to the bar labels in the graphs of Figure 6.

The spike times of all pyramidal cells from all network conditions (from which the SDFs were computed) are available online within the CRCNS repository at http://doi.org/10.6080/K05H7D60 and the calculation of the Spike Density Function is detailed in the Methods section. The power spectra of the SDFs included here were obtained via a one-sided periodogram using Welch’s method where segments have a 50% overlap and a Hamming Window. The highest power within the theta oscillation frequency range of 5 - 10 Hz is reported in Figure 6, and the frequency at which the highest power occurred is reported in Figure 6 - figure supplement 2. Table 6 also lists the power and frequency for each condition.

Figure 6F included three statistically independent simulations from each network condition (control + two experimental conditions). We performed a one-way analysis of variance (ANOVA) of the peak power of the pyramidal cell SDF within the theta frequency range, including all three simulations from each of the three conditions, grouped by condition.

- Pyramidal_SDF_All_Conditions.txt
- Mapping_Network_Condition.txt

